# c-MAF transduces fast motor neuron firing to sustain fast-glycolytic myofibers and neuromuscular junctions

**DOI:** 10.64898/2026.02.05.703983

**Authors:** Edgar Jauliac, Stéphanie Backer, Shunya Sadaki, Julien Gondin, Aurélie Fessard, Hugues Escoffier, Maëva Roullat, Maxime Di Gallo, Adrien Levesque, Doriane Pereira, Matthieu Dos Santos, Vincent Vuong, Alexander S. Ham, Franck Letourneur, Rémi Pierre, Markus A. Rüegg, Carmen Birchmeier, Ryo Fujita, Athanassia Sotiropoulos, Pascal Maire

## Abstract

This study examined how motoneuron activity influences transcription factor binding in mouse fast glycolytic Myh4+ muscle fibers. Single nucleus multiomics of innervated versus denervated tibialis anterior muscles revealed altered chromatin accessibility: SIX and c-MAF binding sites decreased while JUN, FOS, and RUNX1 sites increased in denervated Myh4+ myonuclei. c-MAF showed strong nuclear enrichment after 100 Hz stimulation and periods of increased motoneuron activity but was absent following denervation, establishing it as a primary readout of fast motoneuron firing. Genome-wide analysis demonstrated that c-MAF binding site spacing encodes functionally distinct muscle gene programs. Analysis of constitutive and inducible skeletal muscle-specific c-Maf mutants revealed that c-MAF loss caused region-specific MYH4+ fiber atrophy, MYH1/MYH2 fiber type shifts resembling ALS G93A mouse phenotypes, and progressive neuromuscular junction fragmentation with increased motoneuron terminal sprouting and ectopic reinnervation. These findings establish c-MAF as a critical mediator linking motoneuron activity to muscle gene regulation, fiber integrity, and neuromuscular junction maintenance in fast glycolytic fibers.

## Introduction

Skeletal muscle fibers are innervated by alpha motor neurons which release acetylcholine at slow or fast rates during long or short periods (Kanning et al., 2010). In freely moving adult rats recording from single fast motor units of the fast EDL showed that they were active from 0.5 to 3 minutes/24h, other from 23-72 minutes/24h, while recording from single slow motor units of the slow Soleus showed that they were active from 5.3h-8.4hours/24h (Hennig & Lømo, 1985). Neurotransmitter release at slow 20Hz or fast 100Hz triggers action potentials that propagate along the muscle fiber, initiating calcium release and subsequent myofiber contraction. Calcium is further recaptured by several proteins among which Parvalbumin, Serca1 and Serca2 differentially expressed in slow or fast fibers and allowing fast or slow myofiber relaxation (Schiaffino & Reggiani, 2011)(Gundersen, 2011)(Moreno-Justicia et al., 2025). The resulting calcium peaks also modulate several signaling pathways, with the calcineurin phosphatase pathway in slow oxidative fibers being the most well-characterized. This pathway activates transcription factors such as NFAT and MEF2 (Wu et al., 2000)(Naya et al., 2000)(Olson & Williams, 2000)(Serrano et al., 2001)(Schiaffino & Reggiani, 2011)(Gundersen, 2011) that will activate networks of slow-type/oxidative muscle genes. While mechanisms linking slow motoneuron firing and slow/oxidative muscle gene expression are quite well understood, the signaling cascades that link fast-motoneuron firing and fast-type muscle gene expression in response to increased calcium peaks remain largely unexplored. Furthermore, fast myofibers appear heterogenous: three main subtypes have been characterized according to the *Myosin heavy chain* (*Myh*) gene expressed; MYH being a major constituent of the contractile apparatus. The fast *Myh* (f*Myh*) locus in the mouse is composed of *Myh3, Myh2, Myh1, Myh4, Myh8* and *Myh13* genes arranged within 350Kb on chromosome 11 (Weydert et al., 1985), while the slow *Myh7* gene is located on chromosome 14. Each adult myofiber expresses primarily one *Myh* isoform in all of its myonuclei, underscoring strict transcriptional regulation of fiber identity (Pette & Staron, 2000)(Schiaffino et al., 2015)(Dos Santos et al., 2022), associated with the expression of genes coregulated with each *Myh* gene (Dos Santos et al., 2020)(Petrany et al., 2020)(Kim et al., 2020).

We and others have shown that many genes expressed in adult fast-type myofibers are downregulated upon denervation (Spitz et al., 1998)(Schiaffino & Reggiani, 2011)(Gundersen, 2011)(Collard et al., 2014)(Dyar et al., 2015)(Nakao et al., 2015)(Pillon et al., 2020)(Lin et al., 2022)(Dos Santos et al., 2022)(Lin et al., 2023). Our research has further demonstrated that the SIX/EYA network plays a crucial role in driving fast/glycolytic phenotype (Spitz et al., 1997)(Grifone et al., 2004)(Sakakibara et al., 2014)(Sakakibara et al., 2016)(Wurmser et al., 2020). Single-nucleus Multi-omic (snMulti-omic) experiments have revealed that SIX and MAF transcription factor binding sites are enriched in open chromatin regions specifically within fast/glycolytic *Myh4*+ myonuclei, as compared to other fast-type *Myh1*+ and *Myh2*+ myonuclei (Dos Santos et al., 2020). This finding suggested that MAF transcription factors may act in concert with SIX to generate diversity among fast-type myofiber subtypes. Furthermore, our snMulti-omic data has uncovered distinct gene transcription networks within *Myh4*+ myonuclei across different muscles. This observation potentially links the diversity of skeletal muscles to their anatomical position, specific functions, and the heterogeneity of fast and slow motoneurons along the anteroposterior axis (Dos Santos et al., 2022).

The *Maf* genes encode a subgroup of basic leucine zipper (bZIP) transcription factors that play crucial roles in the regulation of cellular differentiation, tissue development, and homeostasis (Imbratta et al., 2020)(Fujino et al., 2023). They can be divided into two subfamilies: the large MAF proteins (NRL, MAFA, MAFB, and c-MAF), which possess a transcriptional activation domain, and the small MAF proteins (MAFF, MAFG, MAFK), which generally act as transcriptional repressors. Members of this family form homo- or heterodimers, interacting not only among themselves but also with other bZIP transcription factors, including AP-1 family members such as c-FOS, c-JUN, ATF4, and NFE2 (Motohashi et al., 2002). These interactions broaden their regulatory potential and enable context-dependent transcriptional responses. Functionally, MAF proteins may act as pioneer transcription factors, remodeling chromatin to enable subsequent gene activation (Zhu et al., 2018)(Katsarou et al., 2023). Perturbations in *Maf* gene expression or activity have been implicated in tumorigenesis, metabolic disorders, and neurodegenerative or hematological pathologies (Fujino et al., 2023), and *c-Maf* mutant mouse embryos die between E15 to E18 from severe erythropenia (Kusakabe et al., 2011).

More recently Sadaki et al demonstrated the involvement of large MAF proteins in the adult fast subtype phenotype; analysis of triple *MafA/MafB/c-Maf* muscle specific mouse mutants revealed that their absence precluded *Myh4* expression in the fast extensor digitorum longus (EDL), while forced ectopic expression of one of these large MAF in the slow soleus was able to activate *Myh4* (Sadaki et al., 2023). Forced ectopic expression of one of these large MAF in human myotubes was also able to activate *MYH4* (Sadaki et al., 2025)(Dos Santos et al., 2025), a gene expressed mainly in extra ocular muscles in human (L. A. Lee et al., 2019).

Neuromuscular diseases display selective muscle atrophy linked with subtype-specific motor neuron (MN) loss, with fast-fatigable MNs being the most vulnerable in ALS patients and in several mouse models, including the SOD1G93A line (Pun et al., 2006)(Kanning et al., 2010). In amyotrophic lateral sclerosis (ALS), converging evidence supports a multistage pathological process that begins well before overt MN loss, in which cortical and spinal motor circuits exhibit altered excitability and firing patterns and disrupted calcium handling at the neuromuscular junction (Leroy et al., 2014)(Devlin et al., 2015). This functional phase is followed by synaptic retraction and the selective degeneration of fast-fatigable motor units, ultimately leading to denervation and progressive myofiber atrophy (Pun et al., 2006)(Kanning et al., 2010)(Saxena et al., 2013). These findings imply that activity-regulated transcriptional programs within myofibers—particularly those maintaining the fast/glycolytic phenotype—may be disrupted early in disease progression. Furthermore, mutant mice expressing human SOD1 variants specifically in myofibers develop ALS-like phenotypes characterized by oxidative stress, sarcomeric disorganization, mitochondrial dysfunction, and muscle atrophy, even in the absence of widespread MN death, consistent with a primary contribution of muscle in the pathology (Martin & Wong, 2020)(Shefner et al., 2023)(Martínez et al., 2024). These models, together with human SOD1-mutant iPSC-derived myotubes, support the view that disruption of myofiber homeostasis alone can initiate a deleterious “dying-back” process that ultimately impairs motoneuron function and accelerates NMJ degeneration (Badu-Mensah et al., 2020). Understanding how transcription factors such as MAF contribute to preserving muscle identity and neuromuscular junction stability is therefore essential to decipher how motoneuron and muscle dysfunction reinforce each other in ALS.

We show here that c-MAF transcription factor binding observed in myonuclei of *Myh4*+ myofibers was lost upon denervation. We identified dynamic shuttling of c-MAF between the cytoplasm and the nucleus: robust nuclear enrichment of c-MAF was detected during the active period of mice, when motoneuron firing and muscle contraction are increased, as well as after high-frequency (100Hz) NeuroMuscular Electrical Simulation. Furthermore Genome-wide analysis revealed that c-MAF binding sites adopt specific spacing geometries encoding distinct functional programs. Upon denervation, this spacing architecture predetermined chromatin responses, with same-gyre sites undergoing opening to activate developmental reprogramming while cross-gyre and full-nucleosome sites compacted to silence activity-dependent programs. These findings linked fast MN firing to the fast/glycolytic myofiber phenotype. Conditional c-*Maf* ablation in myofibers demonstrated its requirement for the maintenance of *Myh4*+ fibers within distal hindlimb muscles and for NMJ stability, providing a mechanistic link between fast motor neuron activity, fiber-type transcription, and muscle innervation.

## RESULTS

### c-Maf Loss and Motif Switching Drive Transcriptional and Chromatin Remodeling in Denervated Fast Myonuclei

To investigate the role of innervation in the activation and maintenance of the fast-glycolytic program, we generated MULTIOME data (snATAC and snRNA-seq) from *tibialis anterior* (TA) muscle one week after sciatic nerve transection.

Analysis of the data confirmed the presence of the previously described cellular populations in muscle (Figs. 1a, b, S1a). Under denervated conditions, only IIb (*Myh4*+) and IIx (*Myh1*+) myonuclei formed a distinct cluster, whereas denervated IIa (*Myh2*+) nuclei continued to cluster with their innervated counterparts (Supplemental Note1).

**Fig. 1.**
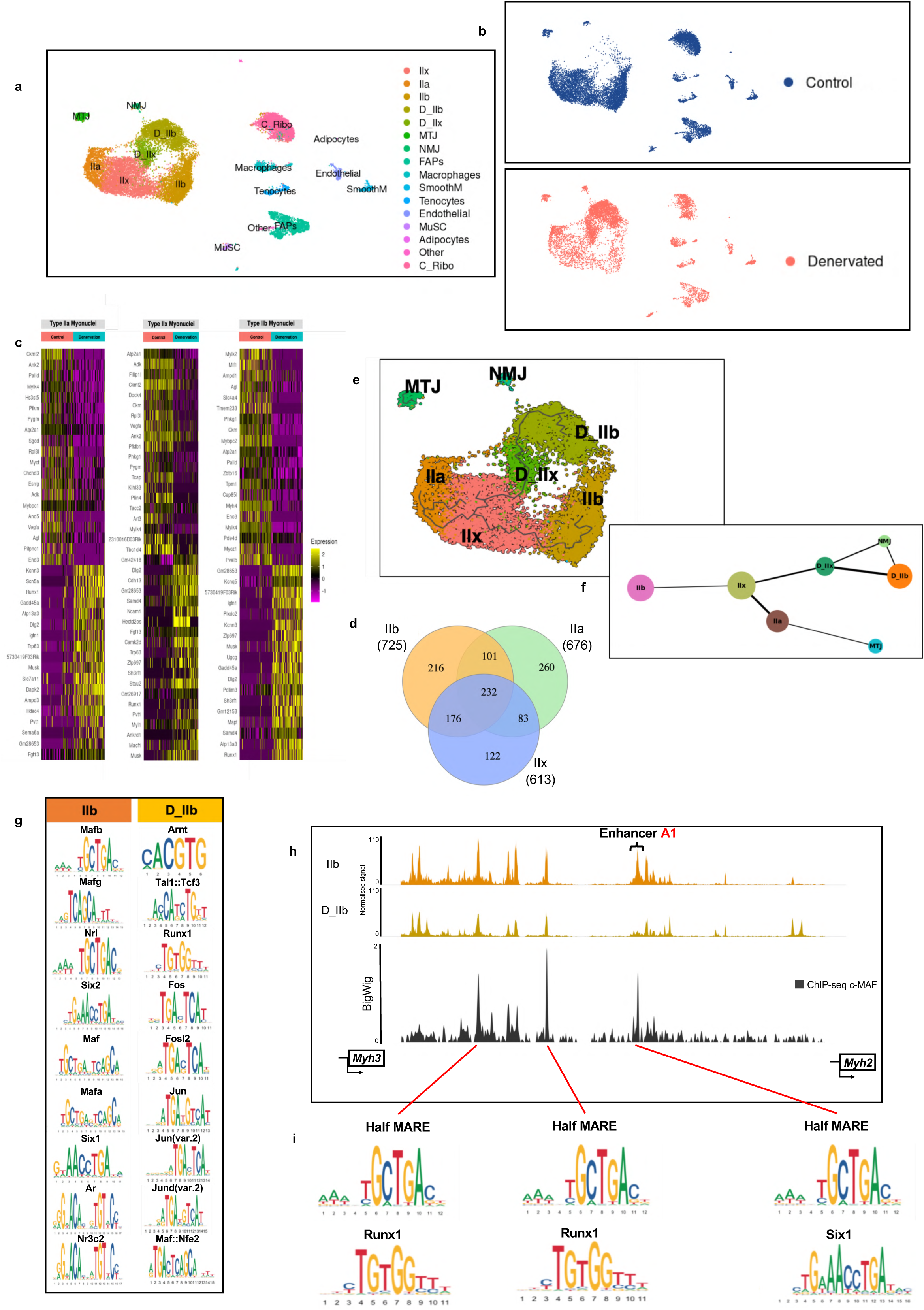
Multi-omic single-nucleus profiling revealed fiber type-specific denervation responses and c-MAF regulatory architecture at the fast *Myosin heavy chain locus*. **a**, UMAP of WNN-integrated snATAC-seq and snRNA-seq from control and 7-day denervated tibialis anterior muscle. Clusters include fiber type myonuclei (IIx, IIa, IIb, D_IIb, D_IIx), specialized populations (NMJ, MTJ), MuSC and non-myogenic cells (FAPs, macrophages, smooth muscle, tenocytes, endothelial, adipocytes). **b**, Split UMAP by condition showing control (blue, top) and denervated (red, bottom) nuclei. Denervated myonuclei partially overlap control clusters. **c**, Heatmaps of top differentially expressed genes in type IIa, IIx, and IIb myonuclei. Color scale: yellow (high), purple (low). Fiber type-specific responses to denervation are visible. **d**, Venn diagram of differentially expressed genes after denervation: IIa/D_IIa (260 unique), IIx/D_IIx (122 unique), IIb/D_IIb (216 unique), with 232 shared across all types. **e**, MONOCLE trajectory analysis showing myonuclei transitions between fiber types and denervation states. Clusters: NMJ, MTJ, D_IIb, D_IIx, IIa, IIx, IIb. **f**, PAGA connectivity graph. Node size indicates cluster size; edge thickness shows connectivity. **g**, Chromvar analysis of Transcription factor binding motif enrichment in type IIb (left column) myonuclei and after denervation (D_IIb, right column). Position weight matrices are displayed. **h**, snATAC-seq coverage at fast *Myosin heavy chain* (f*Myh*) super enhancer. Top track (orange): control, middle track (gold) denervated type IIb accessibility. Bottom track (black): c-MAF ChIP-seq (Dos Santos et al., 2023). Red lines indicate three c-MAF binding regions. **i**, DNA motif composition at c-MAF-bound enhancer regions. Left and middle panels: half-MARE and RUNX1 motifs at distal sites. Right panel: half-MARE and MEF3 (SIX1) motifs at proximal enhancer A1, showing transcription factor architecture.

Differential gene expression analysis revealed significant alterations in the transcriptional programs of IIa/x and IIb fibers (Fig. 1c, S1b, Table S1) (Supplemental Note 2).

PAGA and Monocle3 trajectory analyses (Fig. 1e, f) revealed marked proximity between IIx myonuclei and their denervated equivalents. In contrast, IIb myonuclei showed no direct linkage to denervated counterparts but instead connected with IIx myonuclei. Multiome data demonstrated that *Myh1* expression and chromatin accessibility at its promoter were largely preserved one-week post-denervation (Fig. S1c). Conversely, *Myh2* and *Myh4* genes proved to be more sensitive to denervation, particularly *Myh4*, which displayed ∼90% reduction in both promoter accessibility and transcript levels (Fig. S1c). These findings supported a transient *Myh1*+ state in myonuclei following denervation.

A subset of denervated myonuclei clustered with control IIb myonuclei, indicative of a resilient genetic profile (D_IIb_res, Fig. S1d) characterized by chromatin opening at the *Myh1* promoter with increased transcription, and the opposite for *Myh4* (Fig. S1e). This pattern suggested denervation-induced IIb→IIx fiber type transition, potentially involving loss or downregulation of fast glycolytic regulators including *Six1*, *Nr4a1*, *Tbx15* and *c-Maf* (Chao et al., 2007; Dos Santos et al., 2020; Lin et al., 2022; Sadaki et al., 2023), alongside upregulation of IIx-promoting transcription factors such as AP1 (FOS, JUN) and FOXO family members. Comparative transcriptomic analysis revealed that while both resilient and denervated populations downregulated *Ckm*, *Myh4*, c-*Maf*, and *Nr4a1*, the resilient population displayed a unique signature: selective upregulation of *Myh1*, *Cd36*, and *Foxo1*, plus stable *Tbx15* expression (suppressed in denervated myonuclei), suggesting a protective mechanism against denervation-induced transcriptional reprogramming (Fig. S1f, S2a, Table S3).

We then used Chromvar, a computational tool that infers transcription factor activity variation across single cells or samples by analyzing chromatin accessibility patterns at transcription factor binding motifs.

Chromvar analyses comparing control IIb myonuclei to one-week post-denervation samples revealed loss of characteristic c-MAF, SIX and AR motif enrichment, replaced by RUNX1, ARNT, and AP1 motifs (Fig. 1g, S2a). *c-Maf* expression was markedly reduced following denervation (Fig. S2a, b), paralleling *Myh4* downregulation; many c-MAF target genes overlapped with denervation-altered genes including *Pvalb*, *Atp2a1*, *Actn3* and *Ckm* (Fig. S2c, Table S1).

Upon denervation the 5’-AT-rich MARE half-site (5’-[A/T][A/T][A/T]TGCTGAC-3’) was more robustly suppressed than the canonical palindromic homodimer motif (5’-TGCTGACGTCAGCA-3’). The MAF::NFE2 motif—common to the MAF::AP1 class (5’-[A/G/C]TGA[G/C]TCAGCA-3’, Fig. S2d) was enriched both during denervation and at the neuromuscular junction, suggesting increased MAF::AP1 activity at the NMJ and possible reactivation of the neuromuscular program. During denervation, *c-Maf* was nearly abolished while *Mafg* emerged as the only upregulated *Maf* family member (Fig. S2a), likely associating through AP1 proteins (FOS, JUN, NFE2) or as a negative homodimer.

Super-enhancer mapping at the fast myosin locus (f*Myh*SE) revealed decreased chromatin accessibility, most pronounced in region A1 (Fig. 1h), previously identified as a top peak (Dos Santos et al., 2022). Under denervation, A1 accessibility was nearly abolished (Fig. 1h). ChIP-seq confirmed c-MAF binding to this region, highlighting its innervation- and MAF-dependent regulation. Other c-MAF-binding at f*Myh*SE regions retained greater accessibility under denervation (Fig. 1h). Whereas A1 contains two 5’-AT-rich MARE half-site near a MEF3/SIX binding site, other f*Myh*SE regions harbor MAF::NFE2/AP1 and/or RUNX1 sites, potentially enabling partial accessibility maintenance (Fig. 1i). A1’s accessibility appears uniquely dependent on its sole Half-MARE and SIX binding sites (Fig. 1g, S2b).

To distinguish innervation-dependent from contraction-dependent pathways, we generated single-nucleus RNA-seq from tenotomized *plantaris* muscle, where mechanical unloading occurs without nerve injury. Using Ingenuity Pathway Analysis (IPA), we compared upstream regulators between denervation and tenotomy, identifying those with significant p-values (<0.05) and z-scores (>2 or <−2) exclusive to Myh4+ nuclei versus Myh1+ and Myh2+.

This analysis revealed regulators specifically predicted to be inhibited in denervated *Myh4*+ myonuclei (NOTCH1, KAT2B, MTORC1) or activated (HDAC5, UCP1, SPTBN1) (Fig. S3a-d, Table S4). Notably, HDAC5 is a known repressor of *c-Maf* expression in Schwann cells (Kim et al., 2018) and inhibitor of muscle gene expression (Potthoff et al., 2007), potentially contributing to the observed *c-Maf* suppression following denervation. Furthermore, large MAF transcription factors regulate MTORC1 pathway genes (*Eif4e*, *Deptor*) (Chen et al., 2024), can function as pioneer transcription factors (Zhu et al., 2018), and interact with KAT2B (Bartolome et al., 2018), suggesting that denervation-induced loss of c-MAF may coordinately disrupt both histone acetylation (via KAT2B) and mTORC1 signaling in fast glycolytic fibers.

Interestingly, western blot performed on Cytoplasmic and nuclear extract of control TA showed KAT2B enrichment in tibialis anterior nuclear fractions but near-absence in soleus (Fig. S4a, b).

Furthermore, Luciferase reporter assays revealed synergistic activation between KAT2B and c-MAF on super-enhancer region A1. KAT2B alone could not activate A1, supporting c-MAF recruitment of KAT2B at specific sites. HDAC5 completely repressed A1 activity, which was not rescued by EYA1, or KAT2B. However, c-MAF reversed this repression, restoring baseline activity and supporting its pioneer factor role through half-MARE sites (Fig. S4c, d).

### Nucleosome spacing geometry encodes transcriptional dynamics at c-MAF bound enhancers

The results showing that c-MAF could recruit KAT2B and reverse HDAC5-mediated repression at the A1 super-enhancer raised a fundamental question: how does c-MAF establish and maintain open chromatin configurations at target sites? In eukaryotic genomes, DNA wraps around histone octamers to form nucleosomes, with ∼147 bp of DNA making 1.65 turns around each nucleosome core. The two DNA gyres are parallel to each other, with DNA following a helical path of ∼10 bp per turn. Transcription factor binding sites positioned at specific intervals can both engage nucleosomal DNA: sites within the same gyre are separated by ∼40 bp, while sites on opposing gyres are separated by ∼75 bp but brought into close spatial proximity by the nucleosome wrap. Such positioning enables cooperative interactions on chromatinized DNA without requiring nucleosome eviction. Pioneer transcription factors often exploit this geometry to establish regulatory complexes at compacted loci (Zaret & Carroll, 2011)(Zhu et al., 2018)(Iwafuchi et al., 2020).

The A1 architecture provided a potential answer: two half-MARE sites spaced 37 bp and 77bp apart with an interposed MEF3/SIX1 motif, a configuration compatible with c-MAF binding within the same or cross nucleosomal gyre (Fig 2a). This suggested that nucleosome-compatible spacing might be a genome-wide organizing principle enabling c-MAF to access chromatinized DNA and recruit co-activators like KAT2B. To test this hypothesis, we systematically analyzed half-MARE pair spacing in c-MAF ChIP-seq data (Dos Santos et al., 2023), examining three archetypal nucleosome-compatible configurations: same-gyre spacing (∼40 bp), cross-gyre spacing (∼75 bp), and full nucleosome spacing (∼147 bp). We compared c-MAF in muscle with datasets from other contexts: c-MAF in Th17 lymphocytes (Tanaka et al., 2014), MAFB in macrophages (Soucie et al., 2016), nucleosome binding factor ETV5 in alveolar epithelium (Zhang et al., 2017), KLF4 and SOX2 in MEF, and glucocorticoid receptor (GR) in muscle (Rovito et al., 2021) as a negative control, GR predominantly binds nucleosome-free regions with >95% of sites localizing to pre-existing DNase I hypersensitive sites (John et al., 2011).

**Fig. 2.**
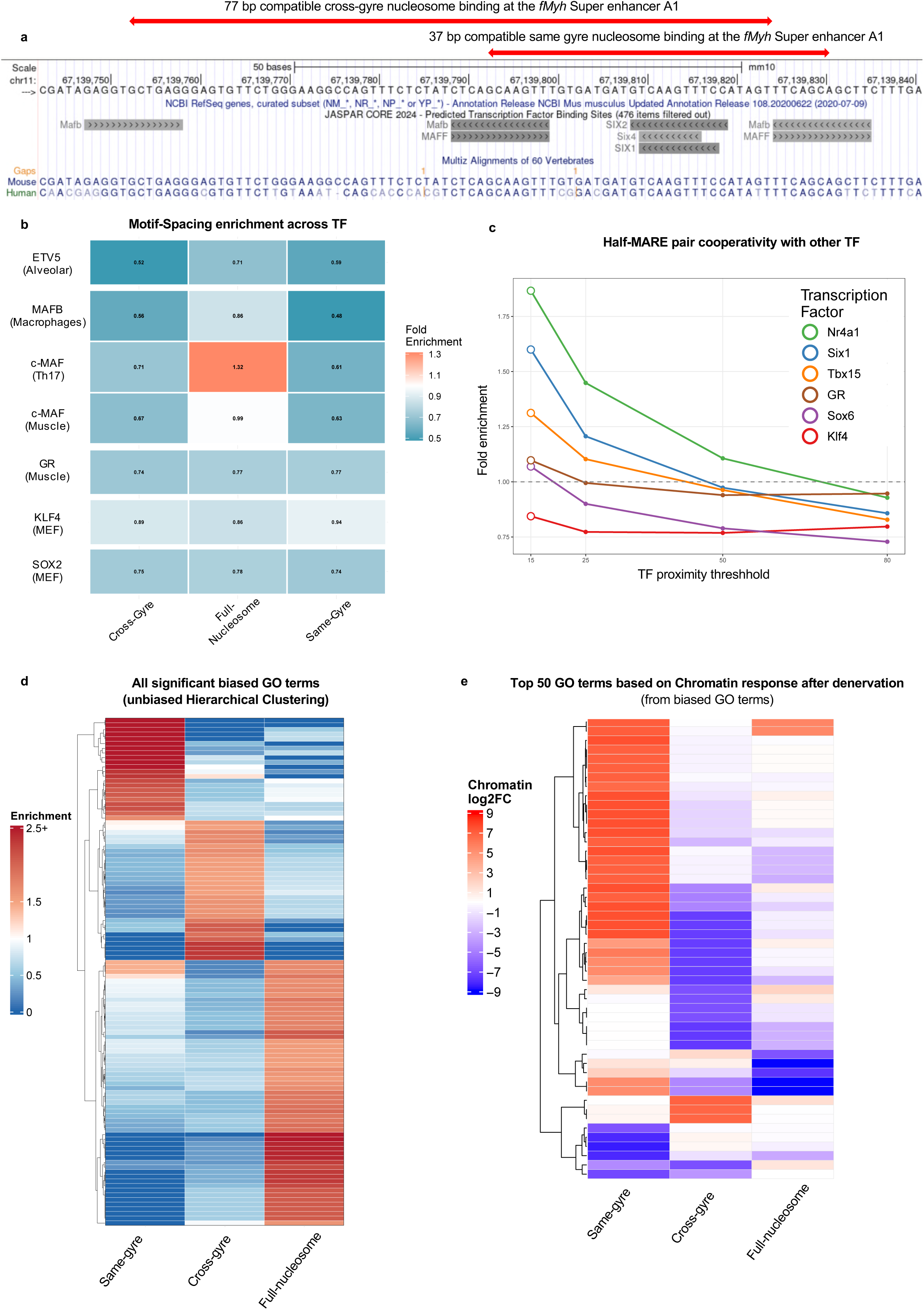
c-MAF exhibits nucleosome-aware chromatin engagement with SIX1-mediated short-range cooperativity. **a**, Sequence alignment of the A1 enhancer of the f*Myh* SE showing 37-bp and 77bp spacing between two conserved half-MARE sites in mouse and human genomes, with a SIX1 binding motif positioned between the two sites. The double red arrow denotes a nucleosome turn. **b**, Heatmap illustrating fold enrichment values (observed/expected) for nucleosome spacing across various transcription factor families and three spacing categories. Color scale highlights depletion (blue) and enrichment (orange), with c-MAF showing relative enrichment at full-nucleosome spacing and depletion at same/cross-gyre categories. **c**, Comparison of cooperativity patterns for SIX1-MARE, TBX15-MARE, NR4A1-MARE, SOX6-MARE, and KLF4-MARE. SIX1, TBX15 and NR4A1-MARE show pronounced short-range cooperativity, while SOX6 and KLF4-MARE show no distance dependence. **d**, Heatmap of all 109 significantly biased GO terms (chi-square p < 0.05) across three nucleosome spacing categories (same-gyre 35–45 bp, cross-gyre 70–85 bp, full-nucleosome 140–155 bp). Rows represent individual GO terms subjected to unbiased hierarchical clustering (Euclidean distance, complete linkage). Color scale indicates enrichment score (observed/expected ratio) for each spacing pattern. Sidebar annotation shows functional category assignment. Full-nucleosome spacing (52.3% of terms, n=57) dominates vesicle transport, protein secretion, and metabolic homeostasis. Cross-gyre spacing (27.5% of terms, n=30) enriches activity-responsive processes including kinase signaling, cell fusion, and migration. Same-gyre spacing (20.2% of terms, n=22) enriches developmental programs including muscle differentiation and morphogenesis. **e,** Heatmap showing chromatin accessibility changes (log₂ fold-change) in denervated IIb myonuclei for the top 50 gene ontology (GO) terms enriched in each c-MAF binding site spacing category. Rows represent individual GO terms, hierarchically clustered by chromatin response pattern. Columns represent the three distinct nucleosome binding geometries.

Analysis revealed that same-gyre and cross-gyre spacings were significantly depleted (observed/expected <1.0) relative to random expectation, while full nucleosome spacing showed no depletion in c-MAF/MAFB datasets (Fig. 2b). Because c-MAF can bind nucleosomal DNA and destabilize nucleosomes (Zhu et al., 2018), this depletion pattern suggests negative selection against configurations that would cause excessive chromatin disruption. Importantly. This depletion pattern was conserved in ETV5 but not in GR, KLF4, or SOX2 datasets, suggesting nucleosome-aware chromatin engagement is a shared property of some nucleosome binding factors.

We next examined cooperativity between half-MARE pairs and other transcription factors. Motif enrichment analysis revealed significant over-representation of NR4A1, SIX1, and TBX15 motifs within 50 bp of half-MARE pairs, with enrichment declining sharply beyond this distance (Fig. 2c). In contrast, KLF4, SOX6, and GR showed no distance-dependent cooperativity, indicating selective co-binding with fast-twitch specification factors. Gene ontology analysis of genes bearing nucleosome-compatible half-MARE pairs revealed c-MAF targets spanning metabolic regulation, NMJ organization, protein secretion, vesicle transport, and ER homeostasis beyond fast-glycolytic genes (Fig. S5a, Tables S5, S6). To identify functional specialization by spacing geometry, we tested whether individual GO terms showed biased representation of same-gyre, cross-gyre, or full-nucleosome configurations among genes with nucleosome-compatible half-MARE pairs. This analysis revealed functional segregation across spacing categories (Fig 2d, Fig. S5b, Table S7). Full nucleosome spacing enriched for vesicle trafficking, ER homeostasis, including COPII components (*Sec24c*, *Sec61g*), ER-to-Golgi transport machinery (*Uso1*, *Cog3*), vesicle targeting Rab GTPases (*Rab4b*, *Rab7b*, *Rab11fip5*). Cross-gyre spacing captured activity-responsive kinase cascades and excitation-contraction coupling machinery: calcium release channels (*Ryr1*, *Cacna1s, Slc8a1, Trdn*), MAP kinase components (*Map2k6*, *Elk1*), PKC isoforms (*Prkce*, *Prkcq*), core contractile proteins (*Myh4* via Enhancer A1, *Mybpc2*, *Actn3*). Same-gyre spacing preferentially associated with developmental transcriptional programs, including master regulators (*Mef2c*, *Smad2*, *Rbpj, Nfic*) and morphogenetic signaling receptors (*Bmpr1a*, *Fzd7*, *Jag1*, *Met*). To test whether c-MAF binding site geometry influences chromatin remodeling, we analyzed accessibility changes in denervated muscle for genes grouped by dominant spacing pattern. Gene ontology terms biased toward same-gyre sites (∼40 bp) showed chromatin opening upon denervation, while cross-gyre (∼75 bp) and full-nucleosome (∼147 bp) sites exhibited chromatin compaction (Fig. 2e, Table S8), probably due to the decreased c-MAF accumulation and absence of binding. Critically, chromatin responses were spacing-specific: within each GO term set, the dominant spacing geometry predicted the strongest chromatin change (Fig. S5c). Motif analysis at chromatin-opening same-gyre sites revealed striking enrichment of HOX homeodomain factors (>130-fold) and AP-1 heterodimers (∼19-fold) (Fig. S5d).

In conclusion, c-MAF binding sites display nucleosome-compatible spacing patterns that segregate functionally distinct gene programs: same-gyre sites regulate repression of developmental program, cross-gyre sites control calcium signaling and contractility, while full-nucleosome sites govern vesicle trafficking and ER homeostasis. Upon denervation, this binding geometry predicts chromatin remodeling responses, with same-gyre sites undergoing chromatin opening at developmental genes, while cross-gyre and full-nucleosome sites undergo compaction.

### Fast Motoneuron Firing Orchestrates c-MAF Nuclear Translocation and Transcriptional Activation in Fast Glycolytic Fibers

Our data, together with published findings (Sadaki et al., 2023)(Dos Santos et al., 2025)(Sadaki et al., 2025), highlighted the central role of c-MAF in the transcriptomic and potentially epigenetic regulation of the fast glycolytic program. We hypothesized that the transcriptional control of c-*Maf* and c-MAF nucleo-cytoplasmic shuttling is driven primarily by motor neuron stimulation rather than by muscle contraction. Following denervation, both the neuron motor signal and muscle contraction are lost, making it essential to disentangle these stimuli.

To investigate the regulation of c-MAF nucleo-cytoplasmic shuttling as a function of activity, we extracted nuclear proteins from the fast TA and from the slow soleus at two circadian time points: ZT03 (Zeitgeber 03, rest phase) and ZT15 (activity phase). During rest phase, nuclear enrichment of c-MAF was higher in the TA than in the soleus (Fig. 3a, S6a), consistent with its heightened role in fast glycolytic fibers that are more abundant in the TA. At ZT15, nuclear enrichment of c-MAF in the TA increased by 70%, whereas no change was observed in the soleus, this suggests that activity and fast motor neuron firing preferentially promoted nuclear translocation of c-MAF in fast glycolytic fibers.

**Fig. 3.**
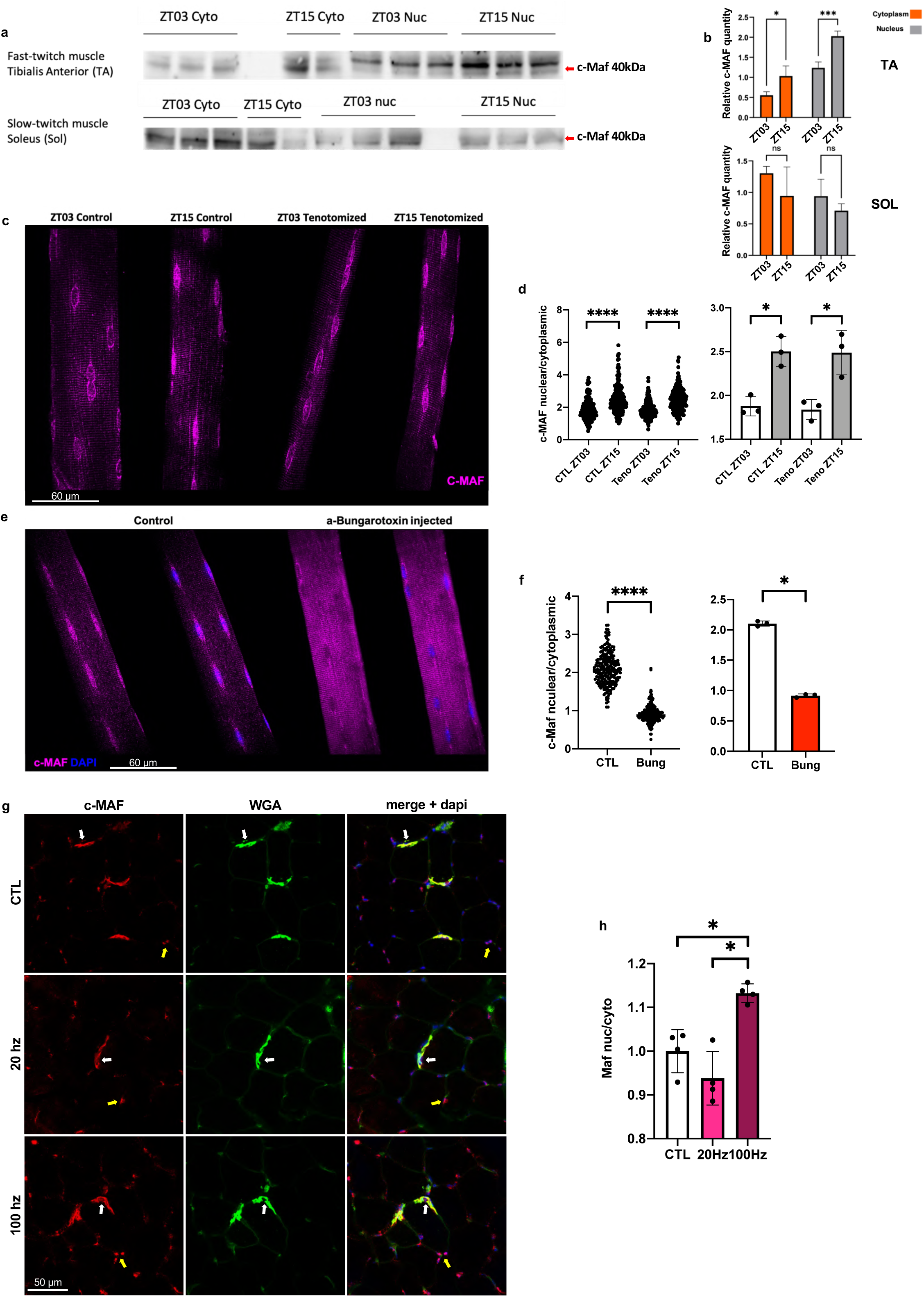
c-MAF nuclear translocation is circadian, activity-dependent, and high frequency-specific in fast-twitch muscle. **a**, Western blot of c-MAF (40 kDa) in cytoplasmic (Cyto) and nuclear (Nuc) fractions from fast tibialis anterior (TA, top) and slow soleus (Sol, bottom) at two circadian timepoints: ZT03 (rest phase) and ZT15 (activity phase). Nuclear c-MAF is enriched at ZT15 in TA; minimal in soleus. **b**, Quantification of cytoplasmic (orange) versus nuclear (gray) c-MAF. TA shows ZT15 nuclear enrichment. Soleus shows lower overall expression with modest circadian variation. Individual data points shown; bars represent mean ± SEM. ***p < 0.001 by two-way ANOVA. **c**, Immunofluorescence of isolated gastrocnemius fibers stained for c-MAF (magenta). At ZT03 control shows modest nuclear signal; ZT15 control displays robust nuclear accumulation. Seven-day tenotomy does not alter nuclear c-MAF accumulation at ZT15. **d**, Plots of c-MAF nuclear/cytoplasmic fluorescence intensity across conditions. On the left plot one dot is one nucleus quantified. The right plot represents average per biological replicates ****p < 0.0001 by one-way ANOVA. **e**, Immunofluorescence of TA isolated fibers at ZT15. Control (left) shows nuclear c-MAF (magenta) with DAPI (blue). α-Bungarotoxin injection (right, blocking neurotransmission) reduced nuclear signal. **f**, Plots showing reduced nuclear/cytoplasmic c-MAF ratio in α-bungarotoxin-treated fibers. On the left plot one dot is one nucleus quantified. The right plot represents average per biological replicates ****p < 0.0001 Unpaired t test. **g**, Confocal images (x60) gastrocnemius cryosections (10 µm) following electrical stimulation. Columns: c-MAF (red), WGA (green, membrane), merge. Rows: control, 20 Hz stimulation, 100 Hz stimulation. Nuclear c-MAF is enriched at 100 Hz, and c-MAF enrichment at NMJ is observed across all conditions. White arrows NMJ. Yellow arrows, myonuclei. **h**, Quantification of nuclear/cytoplasmic c-MAF ratio. 100 Hz stimulation induced significant nuclear translocation compared to control and 20 Hz. *p < 0.05 by one-way ANOVA.

We also observed that the molecular weight of nuclear c-MAF was greater than that of cytosolic c-MAF in the TA but not in the Soleus, suggesting post-translational modifications specific to the nuclear form in the TA such as acetylation or lactylation (Fig. 3a, b).

Although these results identified c-MAF as an activity-responsive regulator, they did not distinguish whether nuclear translocation depends on innervation, contraction, or both. To address this, we performed *gastrocnemius* tenotomy, markedly reducing tension and contractile capacity while preserving innervation, and assessed c-MAF localization by immunofluorescence on isolated fibers at ZT03 and ZT15. In control fibers, nuclear enrichment at ZT15 mirrored western blot results, additionally c-MAF displayed both sarcomeric and perinuclear localization (Fig.3c, d). After tenotomy, nuclear translocation at ZT15 and relative protein abundance were fully maintained, supported by sustain expression according to our single nucleus RNA-seq (Fig. S6b) indicating that c-MAF transcription and localization are directly and exclusively dependent on motor neuron input.

Single-nucleus RNA-seq of muscles treated with botulinum toxin (Ham et al., 2025), a B-toxin family inhibitor of motor neuron neurotransmitter release, revealed almost complete downregulation of *c-Maf*, reproducing the effect of denervation (Fig. S6c). While botulinum toxin is classically known to block the acetylcholine pathway, acetylcholine is not the only neurotransmitter released by motor neurons. Indeed, in Schwann cells, *c-Maf* regulation appears more specifically linked to the NRG1–ERBB2/3/CAMKII pathway rather than Acetylcholine (Ach) dependent pathway (Jo et al., 1995)(Kim et al., 2018).

We then injected α-bungarotoxin into mouse TA muscles at ZT11 on resting mice and gathered the TA muscles at ZT15, to selectively block acetylcholine signaling at the neuromuscular endplate while leaving presynaptic neurotransmitter release intact. Immunofluorescence on isolated fibers revealed that α-bungarotoxin treatment drastically reduced nuclear c-MAF levels while maintaining high transcription and leaving total protein levels seemingly high, suggesting that nuclear translocation of c-MAF depends on neuromuscular activity and associated signaling, rather than on total c-MAF abundance (Fig. 3e, f). This suggested a functional decoupling between transcriptional regulation of *c-Maf*, which seemed activity-independent, and nuclear translocation, which seemed acetylcholine/neural-activity-dependent.

Because c-MAF was strongly enriched and active in nuclei of fast glycolytic fibers, we asked whether its nuclear translocation depends on motor neuron firing frequency. Continuous, low-frequency firing is known to activate the slow oxidative program via the calcineurin/NFAT pathway; however, pathways transmitting the signals of rapid, high-frequency, intermittent firing remain poorly understood (Gundersen, 2011)(Schiaffino & Reggiani, 2011).

We applied patterned electrical stimulation to distal hindlimbs of mice to mimic either fast or slow motor neuron activity (Tothova et al., 2006)(Fessard et al., 2025). Immunostaining for c-MAF on cross-sections revealed that nuclear enrichment occurred specifically after high-frequency stimulation (100 Hz), providing the first evidence that c-MAF relayed rapid motor neuron electrical signals, and identifying c-MAF as the first transcription factor whose nuclear shuttling and nuclear accumulation was under fast motoneuron firing. Yellow arrows indicate c-MAF positive nucleus in all condition with relative enrichment at 100 Hz, white arrows indicate NMJ which appear enriched in c-MAF (Fig. 3g, h, S6d). Additionally, 100 Hz stimulation was not sufficient to induce c-MAF nuclear shuttling in the slow soleus muscle (Fig. S6e). Collectively, these results establish c-MAF as a frequency-dependent sensor that decodes fast motor neuron firing patterns through acetylcholine-mediated nuclear translocation, exclusively in fast glycolytic fibers.

### Loss of c-MAF Recapitulated Regional Muscle Vulnerability Patterns in ALS Through Disrupted Fast Glycolytic Program Control

We next generated both a constitutive muscle-specific *c-Maf* mutant (*cMaf^flox/flox^*;HSA-Cre driven constitutive mutant (Wende et al., 2012)) model and a tamoxifen-inducible mutant model (HSA-CREert2) to better characterize the gene network under the dual control of c-MAF and innervation. As an initial step, we examined the fiber-phenotype of *c-Maf*–deficient mice. Sadaki and colleagues reported that HSA-CRE driven *c-Maf* KO mice showed no shift in fiber-type composition in the EDL or SOL muscles (Sadaki et al., 2023).

Using complete distal hindlimb cross-sections, we observed a clear shift from MYH4+ fibers toward predominantly MYH1+ fibers in the *c-Maf* mutant (Fig. 4a). This shift was region-specific, affecting the peroneal muscles group (PB, PL, FHL, TP, *plantaris*) and the central portion of the TA. In contrast, the EDL displayed no detectable change, as already observed with a distinct *c-Maf^flox^* allele (Sadaki et al., 2023).

**Fig. 4.**
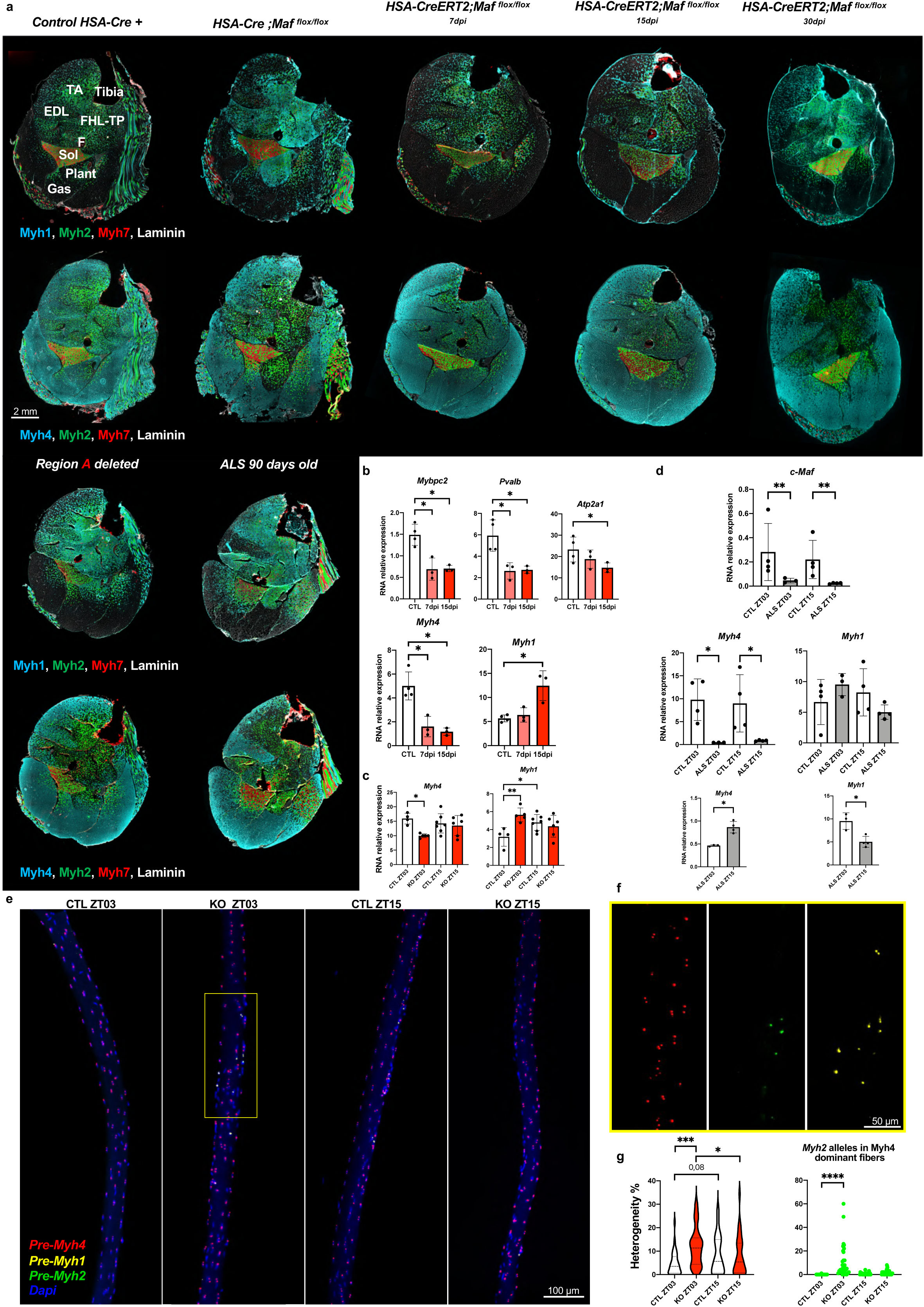
c-MAF loss caused region-specific muscle atrophy with type IIb-to-IIx fiber type transition paralleling ALS pathology. **a**, Immunofluorescence of distal hindlimb cryosections showing MYH isoform distribution and Laminin (white, fiber boundaries). Top rows: MYH1 (cyan, type IIx), MYH2 (green, type IIa), MYH7 (red, type I). Bottom rows: MYH4 (cyan, type IIb), MYH2 (green), MYH7 (red). Columns show: Control *HSA-Cre*+ (leftmost), myofiber constitutive *HSA-Cre;c-Maf^flox/flox^*, myofiber inducible *HSA-CreERT2;c-Maf^flox/flox^* at 7 days, 15 days, and 30 days post-tamoxifen, Region A deleted (f*Myh* super enhancer deletion, *EnhA^-/-^*), and ALS 90 days old. Major muscles labeled in control: TA (tibialis anterior), Tibia, EDL (extensor digitorum longus), FHL-TP (flexor hallucis longus-tibialis posterior), Sol (soleus), Plant (*plantaris*), Gas (gastrocnemius). *c-Maf* loss shows fiber type transition in peroneal group muscles (FHL-TP, central TA) with preserved EDL. ALS models displayed similar regional fiber type switch. **b**, RT-qPCR of fast-glycolytic genes in inducible *c-Maf* mutant 7 and 15 dpi of Tamoxifen. Bottom: *Myh4* and *Myh1* showing temporal changes. *c-Maf* loss reduces *Myh4* expression while delayed *Myh1* increase. Top shows *Pvalb* (parvalbumin), *Atp2a1* (SERCA1), and *Mybpc2* (fast MyBP-C) downregulation. White bars: control (CTL); pink bars: 7 dpi; red bars: 15 dpi. *p < 0.05 by one-way ANOVA with individual data points. **c**, RT-qPCR at 4 weeks post-tamoxifen comparing circadian timepoints ZT03 and ZT15. Left: *Myh4* expression is reduced only at ZT03 (red bars) versus controls (white bars). Right: *Myh1* upregulation in KO at ZT03. *p < 0.05, **p < 0.01 showing sustained fiber type shift. **d**, RT-qPCR on 90 days old ALS G93A mice comparing circadian timepoints ZT03 and ZT15. Top *c-Maf* expression sharply down regulated (grey bars) versus control (white bars). Middle left: *Myh4* expression is constitutively reduced versus controls. Middle right: *Myh1* is upregulated specifically at ZT03. Bottom left *Myh4* expression only in 90 days old ALS mice is upregulated at ZT15. Bottom right same with *Myh1.* **e,** RNA-FISH on isolated TA fibers at ZT03 and ZT15, 4 weeks post-tamoxifen, targeting pre-mRNA of *Myh4* (red), *Myh1* (yellow), *Myh2* (green). DAPI (blue) marks myonuclei. Yellow box shows high-magnification view of transcription foci shown in panel **f**. **g,** *c-Maf* mutant myofibers show increased myonuclei expressing non-dominant *Myh* isoforms, more pronounced at ZT03 than ZT15, indicating spatio-temporal discoordination of the *fMyh* locus. Violin plots, white color control, red *c-Maf* mutant. The % of heterogeneity is the % of f*Myh* allele that differs from the dominant f*Myh* allele per myofiber as estimated by Rnascope experiments. *p < 0.05, ***p < 0.001 ****p < 0.0001 by one-way ANOVA.

This phenotype closely resembled that of mice lacking region A of the f*Myh*SE (Fig. 4a, (Dos Santos et al., 2022), highlighting the strict regulatory control of c-MAF over this enhancer element for the control of the f*Myh* locus. In the inducible mutant, no fiber-type shift was detectable 7 or 15 days after tamoxifen injection (7 or 15 dpi); a pronounced shift was nevertheless observed 4 weeks post-injection (Fig. 4a), suggesting that the changes seen in the constitutive mutant are not attributable to developmental defects.

This MYH shift was accompanied by selective atrophy of resilient MYH4+ fibers in the affected muscles, with no measurable atrophy in the EDL (which instead showed a hypertrophic trend in the constitutive mutant, Fig. S7a, b). Given that c-MAF expression is dysregulated in ALS and our findings implicate c-MAF in fiber-type specification, we asked whether ALS muscles exhibit similar fiber-type transitions (Dos Santos et al., 2025). Strikingly, in ALS G93A model mice, we observed the same MYH4+ to MYH1+ fiber transition pattern, restricted to the same muscle groups and sparing the EDL (Fig. 4a). It also showed the same selective MYH4+ fiber atrophy, with even greater severity. qPCR experiments highlighted rapid downregulation of *Myh4* and fast glycolytic associated genes such as *Mybpc2* and *Pvalb,* on the other hand *Myh1* upregulation is delayed and only apparent at 15dpi (Fig. 4b and c). qPCR results also showed the emergence of circadian regulation of *Myh4* and *Myh1* expression in the *c-Maf* mutant, additionally this temporal change in *Myh4* and *Myh1* expression was also found in 90 days old ALS mice where *c-Maf* is sharply downregulated (Fig. 4c and d), suggesting a role for *c Maf* in maintaining the spatio temporal integrity of the fast-glycolytic program.

Finally, we performed RNAscope on isolated TA fibers at ZT03 and ZT15, targeting the pre mRNA of *Myh4*, *Myh1*, and *Myh2* to assess the potential discoordination of f*Myh* gene expression in *c Maf* mutant TA fibers. *c-Maf* TA mutant myofibers showed increased myonuclear heterogeneity, with more nuclei transcribing f*Myh* isoforms that differed from the fiber’s dominant isoform. This effect was more pronounced during the rest phase (ZT03) than the active phase (ZT15) (Fig. 4e-g). These findings suggest that an additional regulatory factor may partially compensate for the absence of *c Maf* during the active phase, *MafA* and *MafB* being good candidates.

Together, these findings establish that loss of c-MAF recapitulates the region-specific vulnerability pattern observed in ALS through progressive breakdown of spatiotemporal coordination within the fast-glycolytic program, manifesting as increased myonuclear transcriptional heterogeneity and emergence of circadian dysregulation that preferentially impacts MYH4+ fibers.

### c-Maf Deletion uncovered FOXO3-Mediated Atrophy and Spatio-temporal Gene Regulation

Ingenuity Pathway Analysis (IPA) identified upstream regulators driving gene expression changes in *c-Maf* mutant mice and characterized downstream pathological consequences, revealing significant activation of upstream regulators and canonical pathways associated with muscle atrophy, denervation, and neuromuscular dysfunction (Fig. S8a-d) (Supplemental note 3).

IPA predicted that loss of *c-Maf* unleashed FOXO3-driven atrophy coupled with acetylcholine receptor hyperactivation and calcium dysregulation, recapitulating the pathological triad observed in ALS muscle, revealing that c-MAF normally suppresses neurodegeneration-like pathology by directly inhibiting FOXO3-mediated protein degradation programs during denervation (Pikatza-Menoio et al., 2021)(Verma et al., 2022). c-MAF binding intronic enhancer region of *Foxo3* gene reinforced the direct control of *Foxo3* by c-MAF (Fig. S9a).

We validated *in vivo* the activation of FOXO3 by quantifying FOXO3+ nuclei in whole hindlimb sections from *c Maf* mutant mice (Fig. S9b-c). This increase was muscle specific, with marked enrichment in the TA and FHL TP muscles, but no change in the EDL, underscoring the unexpected muscle type specificity of the *c Maf* KO phenotype. Immunofluorescence on isolated TA fibers at both ZT03 and ZT15 revealed persistently elevated nuclear FOXO3 irrespective of circadian timing, with particularly pronounced enrichment in DNA dense nuclear regions at ZT15 (Fig. S9d).

Transcriptomic analysis revealed a disruption of circadian or activity regulation in genes associated with muscle atrophy and protein synthesis (Fig. S9e). *Foxo3*, *Fbxo32*, *Fbxo31*, *Trim63*, *Fkbp5, Prkaa2* and *Ulk2* were chronically upregulated therefore losing their circadian regulation. In contrast, *Map1lc3a* was not constitutively elevated but displayed increased expression exclusively during the rest phase, thereby acquiring circadian regulation. *Myostatin* expression, normally peaking at ZT15, showed exaggerated upregulation at this time point. Then while *Foxo3* was quickly upregulated 7 days after Tamoxifen injection, atrogenes *Fbxo32* and *Trim63* showed delayed upregulation between 7 and 15 dpi (Fig. S9f). Additional alterations were observed in genes linked to protein synthesis and amino acid sensing: *Castor2*, a component of the arginine sensing complex, was chronically upregulated, while the arginine transporter *Slc7a2* and translation regulator *Eif4ebp1* lost their circadian up-regulation at ZT15 (Fig. S9g). Notably, these genes displayed parallel spatio-temporal changes in ALS muscles of 90 days old ALS mice highlighting a subset of spatio-temporal c MAF controlled genes with potential relevance to neurodegenerative muscle pathology (Fig. S9h).

Finally, whole hindlimb immunofluorescence staining for MYH4/2/7 from muscle specific *c Maf* and *Mafb* mutants, and complete *Mafa* mutant mice (Sadaki et al., 2023) demonstrated that the fiber type phenotype and FOXO3/atrogenes regulation were unique to the *c Maf* mutant (Fig. S10a and b). These findings established non redundant transcriptional roles for c MAF, MAFA, and MAFB, and in specific cases such as for the expression of *Eif4e*, even opposing regulatory effects between these MAF family members.

Conversely, multiple anabolic and structural regulators showed significant inhibition in *c-Maf* mutant muscles (Fig. S8b). PI3K/AKT signaling was markedly suppressed, consistent with increased FOXO3 activity. Striated muscle contraction pathways were inhibited, reflecting loss of c-MAF-controlled contractile gene expression (*Actn3*, *Actc1*, *Mybpc2*).

Canonical pathway analysis revealed significant enrichment of neuromuscular pathology pathways (neuropathic pain signaling, acetylcholine receptor pathway). Acetylcholine receptor signaling was significantly activated, indicating potential muscle/motor neuron hyperexcitability reminiscent of ALS pathophysiology (Gunes et al., 2020). Calcium signaling pathways showed robust activation, reflecting disrupted excitation-contraction coupling due to loss of c-MAF-regulated calcium-handling proteins (PVALB, SERCA1). Mitochondrial RNA degradation and rRNA processing pathways were enriched, indicating cellular stress responses and altered protein synthesis capacity. The pathway analysis revealed ALS parallels. Simultaneous activation of acetylcholine receptor signaling, calcium dysregulation, and FOXO3-mediated atrophy mirrors the pathological cascade in ALS muscle (Zhong et al., 2024)(Kubat & Picone, 2024). Corticotropin releasing hormone/dexamethasone activation suggests systemic stress responses extending beyond skeletal muscle, consistent with neuromuscular disease multi-system nature (Fig. S8c). The interplay between altered circadian metabolic regulation and spatio-temporal muscle-nerve coordination is reminiscent of ALS pathophysiology (Fig. S8d; (Huang et al., 2018). This integrated analysis revealed that c-Maf muscle-specific knockout creates a FOXO3-centered pathological network recapitulating neurodegenerative muscle disease features, particularly relevant as c-MAF inhibits FOXO3-mediated atrophy during denervation (Dos Santos et al., 2025) and directly binds a Foxo3 intronic enhancer (Fig S9a).

These findings establish that c-MAF functions as a dual guardian of fast muscle integrity, simultaneously orchestrating spatio-temporal coordination of the glycolytic program while suppressing FOXO3-mediated atrophy. When disrupted, it unleashed a region-specific pathological cascade characterized by loss of circadian metabolic regulation, calcium dysregulation, and acetylcholine receptor hyperactivation, a triad that recapitulates the neuromuscular pathophysiology observed in ALS.

### c-MAF Controled NMJ Architecture Through Activity-Dependent Transcriptional Networks and AchR Stability

Based on the IPA analysis, ALS parallels and the NMJ localization observed in Fig. 3g, we investigated c-MAF’s role at the neuromuscular junction. Immunofluorescence on isolated TA and *gastrocnemius* fibers revealed a pronounced enrichment of c MAF at the NMJ (Fig. S11a). Moreover, approximately 80 genes enriched in subsynaptic nuclei in single nucleus datasets appeared dysregulated in the *c Maf* mutant (Fig. S11b). Among the differentially expressed genes, we noted the upregulation of *Chrnb1*, *Lrp4*, *Kcnq5*, *Musk*, *Sema4d*, *Lrtm1*, while the newly identified NMJ transcript *Pla2g7* was sharply downregulated. Like FOXO3, c MAF binds to the promoter of *Etv5*, a master regulator of the neuromuscular program (Hippenmeyer et al., 2007), via a MARE::AP 1 motif (Fig. S11c). Circadian analysis revealed that *Musk*, *Kcnq5*, and *Pla2g7* expression changes occurred independently of Zeitgeber time, whereas *Lrp4* upregulation was restricted to ZT03, and *Chrna1*, *Runx1*, and *Etv5* upregulation was confined to ZT15. Notably, *Fos* exhibited opposing circadian regulation, being highly upregulated at ZT15 but downregulated at ZT03 (Fig. 5a).

**Fig. 5.**
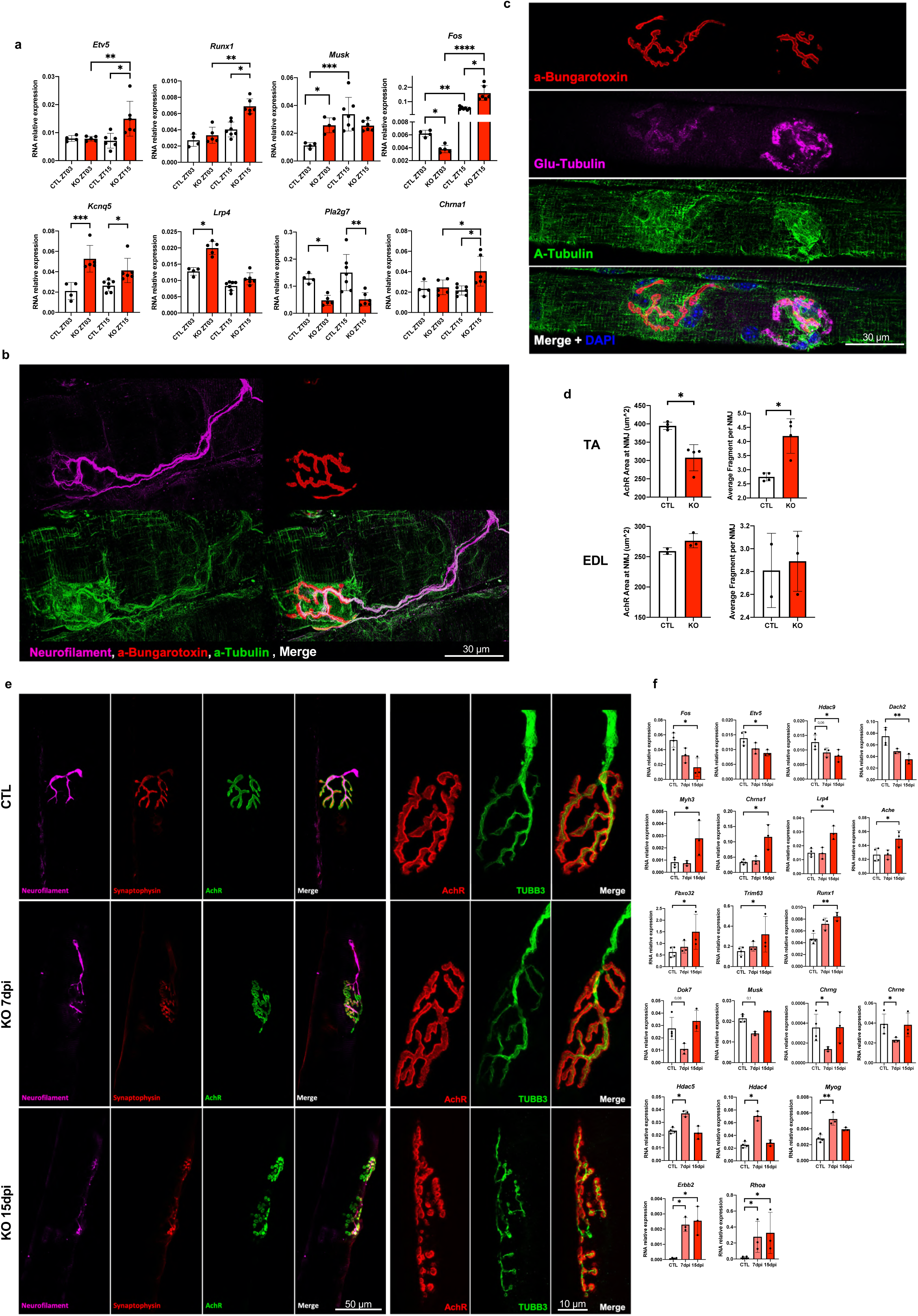
*c-Maf* loss caused progressive NMJ fragmentation with circadian dysregulation of synaptic genes. **a**, RT-qPCR of NMJ-enriched genes at ZT03 and ZT15 4-week post-tamoxifen injection in inducible c-*Maf* mutant. Expression of eight genes is shown: *Etv5*, *Runx1*, *Musk*, *Fos* (top row), *Kcnq5*, *Lrp4*, *Pla2g7*, *Chrne* (bottom row). c-*Maf* mutant disrupts circadian patterns of gene expression. ***p < 0.001, **p < 0.01, *p < 0.05 by two-way ANOVA. **b**, Confocal immunofluorescence of NMJ 4-6 weeks post tamoxifen injection in inducible c-*Maf* mutant. Neurofilament (magenta, presynaptic), α-bungarotoxin (red, AChR), α-tubulin (green, microtubule network) and merge. *c-Maf* mutant NMJ showed fragmented morphology, sign of poly-innervation and axon terminal disorganization. **c**, High-magnification confocal images showing double endplate in *c-Maf* mutant NMJ. Four panels displaying: α-bungarotoxin (red, top), Glu-Tubulin (magenta), α-tubulin (green), and merge with DAPI (blue). Two distinct AChR clusters visible with differential tubulin accumulation patterns on the same mutant myofiber. **d**, Quantification of NMJ morphology in TA and EDL. Left graphs: AChR fragments number per NMJ increased in mutant TA (red bars) versus control (white bars); EDL unchanged. Right graphs: AChR area per NMJ. TA showing significant reduction; EDL preserved. *p < 0.05 (unpaired t-test) demonstrating muscle-specific vulnerability. **e**, Temporal progression of NMJ degeneration at 7- and 15-days post-tamoxifen injection. Three rows (CTL, KO 7dpi, KO 15dpi) with columns showing: Neurofilament (magenta), α-bungarotoxin (red), AChR (green), merge, AChR alone, TUBB3 (green, neuronal Tubulin), merge. Mild destabilization at 7 dpi; extensive at 15 dpi with partial loss of presynaptic innervation. **f**, RT-qPCR panel of 20 NMJ-associated genes comparing CTL (white bars), 7dpi (pink bars), and 15dpi (red bars) of Tamoxifen in *HSA-CreERT2;c-Maf^flox/flox^* TA samples. Coordinated dysregulation indicating systemic NMJ instability and denervation response. *p < 0.05, **p < 0.01, ***p < 0.001 (One-way Anova).

Immunofluorescence staining against AChRs (α bungarotoxin conjugated to a fluorophore) and motoneurons (neurofilament) revealed an increase in fragmented receptor clusters, a reduction in receptor surface area, signs of poly innervation, pre-synaptic cytoskeletal disorganization and more unexpectedly fibers displaying two motor endplates four weeks after tamoxifen injection (Fig.5b-f, S12a-f). The presence of poly innervation and double endplates likely reflects ongoing reinnervation processes, which probably correlated with the observed fiber type shift. These double endplates exhibited features of mature NMJs, notably a characteristic enrichment of the microtubule network (Fig. 5c, S12b-d).

Signs of NMJ instability were clearly detectable and peaked as early as 15 days after tamoxifen administration (Fig. 5e-f, S13a), marked by extensive junctional fragmentation, reduced Synaptophysin coverage at the postsynaptic AChR crests, and partial or complete denervation, neurofilament accumulation and/or absence was observed at the endplate while TUBB3 staining remained more stable, indicative of neuromuscular instability rather than complete denervation (Fig. 5e).

The temporal kinetics of gene expression modulation at 7- and 15-days post-tamoxifen injection (dpi) revealed distinct regulatory categories. First, genes associated with muscle fiber maturation, including *Hdac9*, *Dach2*, *Etv5*, and *Fos*, underwent progressive downregulation, whereas *Runx1* and the aforementioned atrogenes displayed opposite regulation patterns. Second, denervation markers and neuromuscular transcripts, such as *Myh3*, *Chrna1*, *Lrp4*, and *Ache*, showed specific upregulation exclusively at 15 days post-tamoxifen injection.

Interestingly, genes involved in acetylcholine receptor clustering and agrin signaling (*Dok7*, *Musk*, *Chrng*, and *Chrne*) were specifically downregulated at 7 days post-tamoxifen injection, while genes under Neuregulin/CamkII signaling control (*Hdac5*, *Hdac4*, and *Myog*) were upregulated only at this timepoint, displaying a reciprocal expression profile. Finally, *Erbb2* (a neuregulin receptor) and *RhoA* (a downstream target of CaMKII and ERBB2) were sharply upregulated early after Cre-mediated recombination induction (Fig. 5f).

As *Etv5* expression was increased at ZT15 and downregulated 7 and 15 days after tamoxifen injection in *c-Maf* inducible mutant, we performed RNAscope targeting *Etv5* mRNA at ZT03 and ZT15. Globally we observed sharp reduction in *Etv5* puncta number at the NMJ (Fig. 6a). Contrary to qPCR results at ZT15 four weeks after *c-Maf* deletion, *Etv5* appeared markedly downregulated at the NMJ, regardless of circadian timing, with only a modest ZT15 upregulation that failed to restore initial levels. An increased abundance of ectopic *Etv5* transcripts along the myofiber could potentially explain the upregulation observed in qPCR at ZT15 (Fig. 6a-c, S13b-c).

**Fig. 6.**
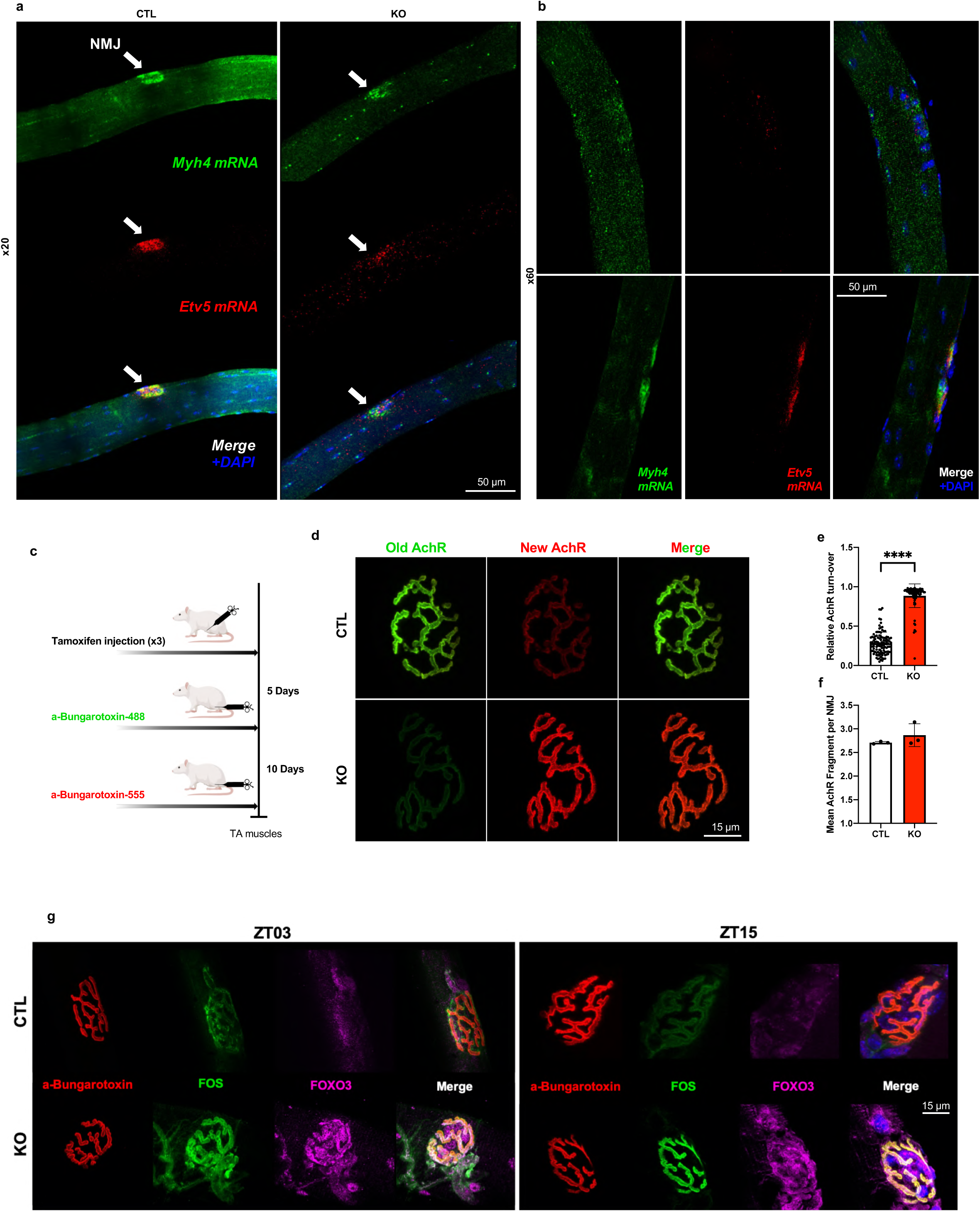
*c-Maf* deletion reduced synaptic *Etv5* expression, accelerates acetylcholine receptor turnover, and induced FOXO3 accumulation at neuromuscular junctions. **a**, Low-magnification (×20) confocal images of RNA-FISH targeting *Myh4* mRNA (green) and *Etv5* mRNA (red) in isolated TA fibers 4-6 weeks post-tamoxifen injection in *HSA-CreERT2;c-Maf^flox/flox^*. Left: Control (CTL) showing abundant *Etv5* expression at NMJ (white arrows). Right: *c-Maf* mutant displayed markedly reduced *Etv5* signal. NMJ regions are indicated by white arrows. **b**, High-magnification (×60) images of *c-Maf* mutant myofibers showing two patterns. Top row: fiber with suspected NMJ location exhibiting almost complete loss of *Etv5* mRNA at NMJ. Bottom row: fiber displaying dual expression pattern with *Myh4* mRNA (green), *Etv5* mRNA (red), and merge. **c,** Experimental timeline and schematic design. *HSA-CreERT2;c-Maf ^flox/flox^* mice received three tamoxifen injections (×3) to induce *c-Maf* deletion, followed by treatment with α-bungarotoxin-488 at 5 days and α-bungarotoxin-555 10 days post-injection to assess AchR turn-over in TA muscles. **d,** Confocal immunofluorescence images of NMJs from control (CTL) and mutant (KO) mice, showing distribution of old acetylcholine receptors (AchR; green) and newly synthesized AchR (red). Representative merged images show overlay of old and new AchR signals. **e,** Quantification of relative AchR turnover in CTL versus mutant mice. Mutant mice show significantly elevated AchR turnover compared to CTL (*** P < 0.001, unpaired t-test). **f,** Quantification of mean AchR fragments per NMJ in CTL and mutant mice. **g,** Confocal immunofluorescence images of NMJs from CTL and mutant mice at circadian timepoints ZT03 and ZT15, showing α-bungarotoxin labeling (red), FOS (green), and FOXO3 (magenta). The merged channel reveals colocalization patterns and temporal dynamics of marker expression. Notably, mutant mice display increased FOXO3 signal and altered FOS/α-bungarotoxin overlap patterns compared to CTL.

We then assessed AChR turnover: we injected α bungarotoxin 488 five days after tamoxifen and a second α bungarotoxin 555 injection 15 days after tamoxifen injection, therefore measuring AChR turn-over on a 10-day timeframe (Fig. 6c), (Falcetta et al., 2024). Isolated TA fibers revealed a drastic increase in 10 day AChR turnover, from an average of ∼25% in controls to nearly 90% in *c Maf* mutant mice, a level characteristic of denervation (Fig. 6d and e). Notably, when AChR receptor activity was inhibited by α-bungarotoxin injection, no fragmentation occurred (Fig. 6f), reinforcing the idea that the “denervation like” phenotype is activity dependent, a relevant observation given that motoneuron hyperexcitability is an early ALS symptom (Bae et al., 2013)(Leroy et al., 2014). We observed a significant increase in c-*Fos* during the active phase (ZT15) (Fig. 5a), and the AP-1 motif is highly enriched in open chromatin of denervated *Myh4*+ myonuclei. These findings led us to consider c-FOS as a potential intrinsic fiber-level marker of denervation. However, c-FOS was not found in nuclei but instead was enriched at NMJ membranes particularly in c-*Maf* mutant mice, where it colocalized with AChRs (Fig. S13d, e). We also detected a marked accumulation of FOXO3 in subsynaptic nuclei and NMJ membranes, potentially explaining part of the increased AchR turnover (Fig. 6g and S13e). Collectively, these findings establish c-MAF as a critical transcriptional regulator of NMJ stability, coordinating the expression of synaptic maintenance genes while preventing activity-dependent AChR destabilization. The activity-dependent nature of the denervation-like phenotype, its prevention by AchR blockade, and the accumulation of c-FOS and FOXO3 at junctional membranes and synaptic nucleus suggest that c-MAF loss triggers a pathological feedback loop where altered transcriptional control and aberrant signaling converge to destabilize the synapse.

Together, these findings reveal that c-MAF functions coordinator of NMJ stability by simultaneously controlling subsynaptic gene expression programs and suppressing activity-dependent AChR destabilization, with its loss increasing FOXO3 in subsynaptic compartments drives accelerated receptor turnover that paradoxically requires ongoing AChR activity, a mechanism directly relevant to ALS pathophysiology.

## Discussion

To identify transcription factors transducing fast motoneuron firing into muscle gene expression programs, we performed chromatin profiling of innervated adult muscle fibers. This revealed specific enrichment of SIX and MAF family binding sites within fast-glycolytic fiber chromatin. We established c-MAF as a master regulator of the fast-glycolytic program, whose nuclear accumulation is controlled by motoneuron firing frequency, the first demonstration of a transcriptional mechanism linking motor neuron electrical activity to fast fiber gene expression in adult muscle.

c-MAF orchestrates fast muscle homeostasis through coordinated regulation of *Myosin heavy chain* genes, suppression of FOXO3-mediated atrophy, circadian metabolic control, and neuromuscular junction stability. c-MAF loss triggered selective fast-fiber atrophy, MYH4-to-MYH1 isoform shifts, and denervation-associated gene activation. Strikingly, these phenotypes recapitulate key features of ALS, where fast motor units show selective vulnerability and preferential degeneration.

The parallels extend beyond fiber-type transitions: both c-MAF deficiency and ALS exhibit region-specific vulnerability (tibialis anterior > EDL), selective fast-fiber atrophy, and progressive neuromuscular junction destabilization. These findings position c-MAF as a critical hub linking motor neuron input to muscle fiber identity and NMJ maintenance. We propose that disruption of the motoneuron-c-MAF-muscle axis represents a fundamental mechanism underlying selective fast motor unit vulnerability in ALS and related motor neuron diseases.

### MAF as a pioneer transcription factor in fast-glycolytic myofibers

Our findings identified c-MAF as the first transcription factor whose nuclear translocation is selectively triggered by 100 Hz high-frequency motor neuron activity, defining a unique transcriptional mechanism tuned to fast motoneuron firing. While NFATC1 couples slow motoneuron activity to gene expression (McCullagh et al., 2004), no equivalent regulator for fast signaling existed until now. Previous work demonstrated that SIX and EYA proteins initiate fast-glycolytic identity (Grifone et al., 2004)(Sakakibara et al., 2014)(Wurmser et al., 2020), with SRF (Charvet et al., 2006) and TBX15 (K. Y. Lee et al., 2015) contributing to its diversity. However, the critical missing element, the transcription factor interpreting fast motoneuron activity, remained elusive.

We show that c-MAF nuclear accumulation is driven by fast motoneuron firing and requires acetylcholine receptor activation. Denervation abolished c-MAF expression and rewired the myonuclear landscape, revealing c-MAF as the pivotal relay mediating fast motoneuron-to-muscle communication. Higher-molecular-weight nuclear c-MAF indicates post-translational modifications, likely acetylation or lactylation. Lactylation is particularly intriguing as it could create a direct metabolic feedback loop linking stimulatory frequency, lactate production, and transcriptional reinforcement of the fast-glycolytic program (Brooks et al., 2023)(Sun et al., 2024).

Emerging evidence suggests c-MAF functions with partial pioneer activity (Zhu et al., 2018)(Katsarou et al., 2023), with position-dependent nucleosome effects. While recent work demonstrated that homotypic motif spacing affects bHLH factor binding strength (Durdu et al., 2025), our analysis demonstrated that nucleosome spacing geometry encodes distinct functional programs. c-MAF employs three DNA spacing categories as a regulatory code: Same-gyre spacing (∼40 bp) enforces stable pioneer binding at developmental regulators (*Mef2c*, *Smad2*, *Rbpj*) insulated from activity modulation. Cross-gyre spacing (∼75 bp) captures activity-responsive modules clustering kinases (*Map2k6*, *PKC*), calcium activators (*Ryr1*, *Cacna1s*), contractile proteins (*Myh4*, *Actn3*). Full-nucleosome spacing (∼147 bp) enables cooperative binding at secretory machinery (*COPII*, *Rab GTPases*) and ER homeostasis. MAF-KAT2B interactions likely recruit SAGA complexes facilitating nucleosomal engagement, with c-MAF overcoming HDAC5 repression to maintain accessibility at f*Myh*SE A1. Denervation responses validated this functional architecture: same-gyre sites undergo chromatin opening and expose HOX/AP-1 binding motifs, activating developmental reprogramming pathways consistent with muscle dedifferentiation. Conversely, cross-gyre and full-nucleosome sites undergo chromatin compaction, silencing activity-dependent contractile machinery and constitutive secretory programs during atrophy. This spacing-specific chromatin remodeling demonstrates that nucleosome geometry predetermines transcriptional responses to physiological perturbation, with c-MAF binding site architecture encoding both steady-state fiber identity and denervation-responsive programs.

### Loss of c-MAF Unmasks HDAC4/5-Mediated Repression and Atrophy Pathways in Denervated Fast Muscles

Our findings resolved a longstanding paradox: how HDAC4/5, enriched in fast fibers and repressing MEF2-dependent transcription (Potthoff et al., 2007)(Cohen et al., 2015), permitted robust fast-glycolytic gene expression under normal innervation yet contributed to suppression following denervation. We demonstrated that c-MAF, robustly expressed under innervation and downregulated following denervation, actively displaced HDAC4/5 from fast-glycolytic enhancers. Through luciferase assays, we showed c-MAF uniquely reversed HDAC5-mediated repression of f*Myh* super-enhancer A1, whereas SIX1, EYA1, and KAT2B could not. This positioned c-MAF as an innervation-dependent guardian of fast-fiber identity counterbalancing HDAC4/5 repressive potential (Potthoff et al., 2007). Upon denervation, c-MAF loss allowed HDAC4/5 reoccupation of fast-gene regulatory elements, converting them to repressed states and driving atrophy and fiber-type destabilization. c-MAF loss triggered metabolic reprogramming through FOXO3-centered atrophy network activation (Sartori et al., 2021). Loss of *c-Maf* caused circadian disruption, chronically dysregulating the protein synthesis-proteolysis balance critical for muscle homeostasis. Notably, FOXO3 activation showed regional specificity, with pronounced nuclear enrichment in *tibialis anterior* and *flexor hallucis longus/tibialis posterior* muscles while sparing the *extensor digitorum longus*. This pattern correlated with fiber-type transition dynamics, suggesting muscles with greater fiber-type heterogeneity and stronger c-MAF reliance are particularly vulnerable. The fast/glycolytic MYH4+ fibers of the EDL, rarely engaged during routine cage activity (Hennig & Lømo, 1985), may sustain their identity independently of continuous motoneuron input. In contrast, larger MYH4+ fibers of the TA, activated more frequently, depend heavily on sustained motoneuron-derived signals. Furthermore, MAFA and MAFB may provide varying compensation, as triple mutants show reduced MYH4 myofibers in EDL (Sadaki et al., 2023). We cannot rule out that distinct developmental trajectories between the EDL and TA muscles, shaped by lineage-specific regulatory landscapes established during limb morphogenesis, may contribute to the differential dependence of their fast MYH4+ fiber characteristics on motoneuron firing activity. The molecular heterogeneity observed among *Myh4+* myonuclei in adult limb muscles (Dos Santos et al., 2020) could thus reflect the persistence of distinct embryonic trajectories influencing adult muscle phenotype and neuromuscular responsiveness.

### Transcriptional Control of Synaptic Homeostasis by c-MAF Connects NMJ Integrity to Neurodegenerative Susceptibility

c-MAF loss triggered neuromuscular junction instability characterized by fragmentation, poly-innervation, and double endplate formation. AChR turnover accelerated from ∼25% to 90% in *c-Maf* mutants, reaching denervation-characteristic levels and reflecting fundamental postsynaptic apparatus destabilization, as already observed in ALS mouse models (Vinsant et al., 2013)(Tremblay et al., 2017). Critically, α-bungarotoxin addition completely prevented NMJ breakdown, demonstrating that instability is activity-dependent and directly linking motor neuron excitability to progressive synaptic deterioration. c-MAF loss integration with ALS pathophysiology revealed phenotypic parallels. Identical regional fiber transitions (MYH4→MYH1/2 shifts) occurred in c-*Maf* mutant and ALS G93A models with remarkable specificity affecting peroneal muscle groups while sparing EDL. This selective vulnerability may reflect differential fast fiber atrophy patterns and intrinsic collateral reinnervation capacity (Ovsepian et al., 2023). Selective atrophy predominantly affects non-transitioning MYH4+ fibers at muscle peripheries with homogeneous fiber-type composition, suggesting lower collateral reinnervation probability. Regions undergoing transition are in closer proximity to slower motor units due to heterogeneous fiber-type composition. Temporal progression studies demonstrate that NMJ changes preceded motor neuron death in multiple ALS models, supporting the "dying back" hypothesis (Martin & Wong, 2020)(Shefner et al., 2023)(Martínez et al., 2024) while positioning c-MAF-dependent pathways as early pathological events. The link between early ALS and the activity-dependent *c-Maf* mutant phenotype establishes c-MAF as a critical node whose activity-dependent regulation connects neuromuscular homeostasis to neurodegenerative vulnerability.

## Acknowledgements

We thank G.Macaux for meaningful discussion regarding bio-informatic analysis, F.Britto, L.Giordani, J.Smith, V.Gache, P.Castets and S. Schiaffino for meaningful scientific discussion. We thank Emma Castagnet and Marcio Do-Cruzeiro of the MOUST’IC core facility and Antoine Guéraud of the Institute Cochin for their technical assistance. We thank J.S.Annicotte, C.Pouponnot and S.Emiliani for the gift of Kat2b, c-Maf and Hdac4 expression vectors respectively. E.J is supported by a PhD fellowship from the University Paris Cité and from the Association française contre les myopathies (AFM). Financial support was provided by the IdEx Université Paris Cité (ANR-18-IDEX-0001-NEUROMYO), AFM MYHREG (n°23012), AMM (MYOREPRO), ANR MOTOMYO (ANR-21-CE14-0042-01), the « Fondation pour la Recherche Médicale » (FRM, EQU202503020009), the Institut National de la Santé et de la Recherche Médicale (INSERM), the Centre National de la Recherche Scientifique (CNRS).

## Author Contributions Statement

Designed experiments: E.J., A.S., S.B., M.D.S, J.G, P.M., Performed experiments: E.J., S.S., D.P., J.G., A.S., A.F., H.E., M.R., M.D.G., A.L., V.V., F.L., M.D.S., S.B., D.J.H., R.P. Interpreted the data : E.J., A.S., S.B., S.S., C.B., M.R., R.F., P.M. Wrote the manuscript : E.J., P.M.

## Supplemental Notes

**Supp 1.** We also noted the presence of a novel cluster named C-Ribo enriched in *Gm26917*, *Gm42418 and Malat1* transcripts described previously as ribosomal contamination in single-cell studies, here we used single-nucleus and the biological relevance of this cluster remained unclear, we did not include it in subsequent analysis (Figs. 1a, b).

**Supp 2.** The expression of 232 genes was commonly modulated in *Myh4* (IIb), *Myh1*(IIx) and *Myh2* (IIa) *+* myonuclei, while the expression of 216, 122 and 260 genes was modulated only in *Myh4*+, *Myh1*+ and *Myh2*+ myonuclei respectively (Fig. 1d, Table S2). We observed in all types of myonuclei the down regulation of *Atp2a1* and *Mylk4* and the upregulation of *Runx1, Dlg2* and *Musk*, associated respectively with the atrophy response and neuromuscular program. Among the genes specifically modulated within fiber subtypes, we noted the down regulation of *Maf*, *Pvalb* and *Tbx15* and the up regulation of *Kcnk5* and *Atp1a2* in *Myh4*+ myonuclei, the down regulation of *Pbx1* and *Lpl* and the up regulation of *Myh10, Sntb1* and *Ldlrad3* in *Myh2*+ myonuclei.

**Supp 3.** Upstream regulator analysis identified FOXO3 as the most significantly activated transcriptional regulator, consistent with its master regulator role in muscle atrophy. Dexamethasone showed predicted activation, indicating glucocorticoid stress response amplifying FOXO3-mediated protein degradation. The metabolic stress sensor AMPK was activated, reflecting impaired muscle energy homeostasis. Additional activated regulators (PLCG2, PRKCD, PPP2CA) indicated dysregulated calcium signaling and kinase-phosphatase networks linked to FOXO3 regulation (Fig. S8a-d). Upstream regulator analysis (Fig. S8e) identified FOXO1 as a prominently activated master regulator of muscle atrophy in both *c-Maf* mutant and denervated muscle, alongside metabolic stress sensors (TXNIP, LDHA) and inflammatory/stress-response regulators (CREB1, TNF, PRKCD), while muscle maintenance factors (DMD, COLQ, IL15) were suppressed. Canonical pathway analysis (Fig. S8f) revealed widespread activation of neuromuscular junction-related signaling (acetylcholine receptor, synaptogenesis, calcium signaling), mitochondrial dysfunction, and NRF2-mediated oxidative stress responses, accompanied by profound inhibition of glycolytic and gluconeogenic pathways. This molecular signature, characterized by FOXO-driven catabolism, AMPK-mediated metabolic stress, and dysregulated calcium/kinase networks, demonstrated that loss of c-MAF recapitulates the transcriptional hallmarks of denervation-induced muscle atrophy and neuromuscular junction destabilization. The dysregulation of chromatin organization pathways (Fig. S8f) in *c-Maf* mutant muscle provides critical mechanistic insight into c-MAF’s function as a potential pioneer transcription factor capable of accessing closed chromatin. Earlier nucleosome spacing analysis demonstrated that c-MAF binding sites exhibit distinctive nucleosome positioning patterns, suggesting c-MAF actively remodels chromatin architecture to establish permissive transcriptional states for muscle fiber identity genes. The profound disruption of chromatin organization upon c-MAF loss, coupled with dysregulation of calcium signaling and neuromuscular junction pathways, reveals a mechanistic link between transcriptional pioneer activity and neuromuscular stability. This connection may be particularly relevant to understanding shared pathogenic mechanisms in demyelinating and motor neuron diseases: multiple sclerosis involves oligodendrocyte dysfunction and myelin loss leading to secondary axonal degeneration, while ALS exhibits primary motor neuron degeneration with subsequent muscle denervation. Both diseases converge on neuromuscular junction destabilization and muscle atrophy, processes that our data suggest require intact c-MAF-mediated chromatin remodeling to maintain the transcriptional programs linking neuronal activity to muscle fiber homeostasis. The loss of pioneer factor activity at critical regulatory elements may thus represent a common vulnerability in neurodegenerative cascades affecting motor systems (Fig. S8e,f).

## SUPPLEMENTARY TABLES

**Supplementary Table S1. Differential gene expression analysis in denervated versus control myonuclei across fiber types.**

Single-nucleus RNA-sequencing differential expression analysis comparing denervated and control conditions within fiber type-specific myonuclei populations (Type IIb, Type IIx, and Type IIa). For each fiber type, the table contains: gene symbol, p-value (two-sided Wilcoxon rank-sum test), average log2 fold-change (denervation vs. control), percentage of nuclei expressing the gene in denervation condition (pct.1), percentage of nuclei expressing the gene in control condition (pct.2), and Bonferroni-corrected p-value (p_val_adj).

**Supplementary Table S2. Venn diagram analysis of denervation-responsive genes across muscle fiber types.**

Gene lists representing the intersection and fiber type-specific differentially expressed genes following denervation in Type IIb, Type IIx, and Type IIa myonuclei. Genes were categorized into seven groups based on their differential expression pattern: (1) IIb ∩ IIa ∩ IIx (232 genes): differentially expressed in all three fiber types, representing a core denervation response program; (2) IIb ∩ IIx (176 genes): shared response between fast-glycolytic fiber types; (3) IIb ∩ IIa (101 genes): shared response between Type IIb and IIa; (4) IIb-specific (216 genes): uniquely responsive in Type IIb myonuclei; (5) IIa-specific (260 genes): uniquely responsive in Type IIa myonuclei; (6) IIa ∩ IIx (83 genes): shared response between Type IIx and IIa; (7) IIx-specific (122 genes): uniquely responsive in Type IIx myonuclei.

**Supplementary Table S3. Venn diagram analysis comparing denervation-responsive genes in resilient versus classic Type IIb myonuclei.**

Gene lists representing the differential expression response to denervation in two functionally distinct Type IIb myonuclei subpopulations. Genes were categorized into three groups based on their differential expression pattern (vs. control innervated IIb): (1) IIb_D_res ∩ IIb_D (268 genes): differentially expressed in both resilient and classic denervated IIb myonuclei, (2) IIb_D_res-specific (37 genes): uniquely differentially expressed in resilient denervated IIb myonuclei, (3) IIb_D-specific (457 genes): uniquely differentially expressed in classic denervated IIb myonuclei,

**Supplementary Table S4. Upstream regulator analysis of differentially expressed genes in fiber type-specific myonuclei following denervation or tenotomy.**

Ingenuity Pathway Analysis (IPA) upstream regulator analysis identified transcriptional regulators, signaling molecules, and other factors predicted to be activated or inhibited based on differential gene expression patterns in Type IIb, Type IIx, and Type IIa myonuclei following denervation (Dener) or tenotomy (Teno) compared to control conditions. The table contains 3,109 putative upstream regulators with their predicted activation state represented as z-scores for each fiber type and perturbation condition. Positive z-scores indicate predicted activation of the upstream regulator, while negative z-scores indicate predicted inhibition. (z-score threshold used was >2 or <-2). Notably, 45 upstream regulators are specifically predicted in IIb denervation but not in IIx or IIa denervation nor tenotomia, including HDAC5 (z-score = 2.412, predicted activated) and KAT2B (z-score = -2.646, predicted inhibited), indicating fiber type-specific regulatory responses to denervation. Z-scores were calculated by IPA based on the overlap between observed differential gene expression patterns and known downstream targets of each upstream regulator, with enrichment for genes changing in a direction consistent with regulator activation or inhibition.

**Supplementary Table S5. Venn diagram analysis of genes with c-MAF binding sites categorized by half-MARE pair spacing geometry.**

Gene lists representing the spatial organization of c-MAF binding sites based on half-MARE pair spacing patterns identified by ChIP-seq analysis. Half-MARE pairs were classified into three geometric categories: (1) Same-Gyre (956 genes total) (2) Cross-Gyre (1,156 genes total) (3) Full-Nucleosome (1,052 genes total).

**Supplementary Table S6. Gene Ontology enrichment analysis of c-MAF target genes harboring half-MARE pair.**

Gene Ontology (GO) enrichment analysis of genes containing c-MAF half-MARE pair. The table contains 1,433 significantly enriched GO terms (q-value < 0.05) categorized into Biological Process (BP, n=1,163), Molecular Function (MF, n=143), and Cellular Component (CC, n=127). For each GO term, genes were analyzed for the spatial organization of their c-MAF binding sites across three geometric configurations: Same-Gyre, Cross-Gyre and Full-Nucleosome. The table provides: GO term ID and description, total gene count per term (Count), number of genes in each spacing category (n_same, n_cross, n_full, n_none), percentage distribution across spacing patterns (pct_same, pct_cross, pct_full, pct_none), enrichment scores for each spacing pattern relative to background distribution (enrichment_same, enrichment_cross, enrichment_full), statistical significance of spacing pattern bias (chi-square and Fisher’s exact test p-values), GO term enrichment significance (go_pvalue, go_qvalue), and functional categorization.

**Supplementary Table S7. Gene Ontology terms with significant chromatin spacing bias for c-MAF binding sites identified by chi-square analysis.**

Subset of 109 GO terms from Supplementary Table 6 displaying significant non-random distribution of c-MAF half-MARE pair spacing patterns (chi-square test, p < 0.05 for at least one comparison). These terms represent biological processes, molecular functions, and cellular components whose c-MAF-regulated genes preferentially utilize specific chromatin binding geometries. GO terms are categorized into Biological Process (BP, n=92), Molecular Function (MF, n=11), and Cellular Component (CC, n=6). For each biased GO term, the table provides: GO term ID and description, total gene count per term (Count), distribution of genes across spacing patterns (n_same, n_cross, n_full, n_none for Same-Gyre, Cross-Gyre, Full-Nucleosome, and no-defined-spacing respectively), percentage representation (pct_same, pct_cross, pct_full, pct_none), enrichment scores relative to background distribution (enrichment_same, enrichment_cross, enrichment_full), the predominant spacing pattern (dominant_spacing), statistical significance measures (chi-square p-value, Fisher’s exact test p-values for each spacing category), GO enrichment statistics (go_pvalue, go_qvalue), and functional annotation. The dominant spacing pattern distribution reveals 57 GO terms enriched for Full-Nucleosome spacing, 30 for Cross-Gyre spacing, and 22 for Same-Gyre spacing.

**Supplementary Table S8. Gene Ontology terms enriched in c-MAF binding sites show spacing-specific chromatin responses to denervation**

Gene Ontology (GO) biological process terms significantly enriched at c-MAF binding sites in Myh4+ myonuclei (n = 1,392 terms), categorized by dominant c-MAF binding site spacing pattern and chromatin accessibility response upon denervation. Dominant_Spacing indicates the spacing category (same-gyre ∼40 bp, cross-gyre ∼75 bp, or full-nucleosome ∼147 bp) showing strongest c-MAF binding site enrichment for each GO term. n_genes and n_regions_total denote the number of genes and total c-MAF binding regions associated with each term, respectively. same_gyre, cross_gyre, and full_nucleosome columns report mean log fold change in chromatin accessibility (denervated versus innervated Myh4+ myonuclei) at c-MAF binding sites within each spacing category for the given GO term; positive values indicate increased accessibility (chromatin opening), negative values indicate compaction. n_same_gyre, n_cross_gyre, and n_full_nucleosome indicate the number of c-MAF binding regions in each spacing category for the respective GO term. GO terms demonstrate spacing-specific chromatin remodeling during denervation-induced muscle atrophy: contractile machinery and metabolic genes (cross-gyre dominant) undergo chromatin compaction, whereas developmental regulators (same-gyre dominant) exhibit chromatin opening and secretory/signaling pathways (full-nucleosome dominant) show category-specific responses. This spacing-dependent chromatin behavior validates that nucleosome geometry encodes both fiber-type identity programs and denervation-responsive transcriptional circuits.

## MATERIELS and METHODS

Animal experimentation adhered strictly to the institutional guidelines for care and use of laboratory animals as outlined in the European Convention STE 123 and the French National Charter on the Ethics of Animal Experimentation (n°44083-2022121514506347, n°2024022718312758 and n°201809041512432, agreement n°C75-14-02). Animal procedures received approval from the French Ethical Committee of Animal Experiments CEEA - 034 and were performed in Institut Cochin animal core facility (Agreement A751402).

### Tissue Preparation and Processing

#### Tissue Collection

Fast-twitch (Tibialis Anterior, TA) and slow-twitch (Soleus, Sol) muscles were dissected from 3 months old mice at ZT03 and ZT15 timepoints and immediately processed for subcellular fractionation.

#### Surgical Interventions

For tenotomy experiments, the distal tendon of the gastrocnemius muscle was surgically severed under isoflurane anesthesia. For α-bungarotoxin injection experiments, tibialis anterior (TA) muscles were injected with 10 µg/mL of α-bungarotoxin (Thermo Fisher Scientific) dissolved in sterile PBS. Control muscles received equivalent volumes of PBS. For denervation experiments, sciatic nerve transection was performed on 8-week-old male and female mice. Muscles were harvested at ZT03 and ZT15 timepoints or X days post-injection (α-bungarotoxin), 7 days post-surgery (denervation and tenotomy), and immediately fixed in 4% paraformaldehyde (PFA) in PBS for 20-30 minutes at +4°C or room temperature after +37°C pre-heat for Tubulin staining, then washed three times in PBS.

#### Tissue Processing and Cryosection

Entire hindlimbs were dissected and embedded in optimal cutting temperature (OCT) compound, then frozen in liquid nitrogen-cooled isopentane. Transverse cryosections (10 µm) were cut using a cryostat (CM3050 S model, LEICA manufacturer) and mounted on Superfrost Plus glass slides (Thermo Fisher Scientific). Sections were stored at -80°C until use.

#### Single Muscle Fiber Isolation

Individual muscle fibers were mechanically isolated from fixed gastrocnemius and TA muscles by microdissection in PBS containing 0.05% Triton X-100 using fine-tipped tweezers. For *in vivo* AChR labeling experiments, hindlimbs were fixed intact (muscle on bone) in 4% paraformaldehyde (PFA) in PBS for 20 minutes at room temperature on a shaker, protected from light. Following fixation, tissues were rinsed twice with PBS containing 0.1 M glycine for 10 minutes each wash and stored at 4°C if necessary. Myofibers were then isolated as described above.

#### Cell culture and transfection experiments

All cultures were grown in humidified incubators at 37°C under 5% CO_2_. Myogenic primary cells were seeded on 1% Matrigel in growth medium (GM) containing DMEM/F12, 20% FBS, 1X Ultroser™ G and 1X antibiotic-antimycotic (anti-anti). For differentiation, myoblasts were seeded in Matrigel-coated dishes and cultured in differentiation medium (DM) (DMEM/F12, 2% Horse Serum, 1X anti-anti). Myogenic primary cells were seeded on 5% Matrigel-coated dishes in growth medium (GM) containing DMEM/F12, 20% FBS, 2% Ultroser™ G and 1X antibiotic-antimycotic (anti-anti). For differentiation, myoblasts were seeded in Matrigel-coated dishes and cultured in differentiation medium (DM) (DMEM/F12, 2% Horse Serum, 1X anti-anti). After 3 days in proliferation medium, cells are transfected using Lipofectamine 3000 (Invitrogen). The Enhancer A1 region was amplified by PCR from mouse genomic DNA using the following primers: forward 5’-TTTGACGTCGCTAACTATCGGAATGTAA-3’ and reverse 5’-AAAACTGCAGCGATTGATAGCCTTACATT-3’. One hundred nanograms of expression vector are added to 800 ng of the EnhA1-Luciferase vector and 10ng of tk-Renilla vector in a mixture containing 1 microgram of DNA. The DNA–Lipofectamine complex is applied to 100,000 cells that have been in differentiation medium for 24 hours in a well of 1.9 cm². Twenty-four hours after transfection, the transfected myotubes are lysed with passive lysis buffer in the Dual Glo Luciferase assays (Promega). Luciferase and Renilla activities were measured with the Centro LB960 Luminoskan plate reader. The results shown are normalized by tk-Renilla activity.

### Protein Analysis

#### Nuclear and Cytoplasmic Protein Extraction

Nuclear and cytoplasmic protein fractions were prepared using the NE-PER Nuclear and Cytoplasmic Extraction Reagent kit (Thermo Fisher Scientific) according to the manufacturer’s instructions. Briefly, frozen muscle tissue was homogenized in ice-cold CER I buffer supplemented with protease and phosphatase inhibitor cocktail (1:100, Roche). Following a brief incubation on ice, CER II buffer was added to facilitate cytoplasmic protein extraction. Samples were vortexed and centrifuged at 16,000 × g for 5 minutes at +4°C. The supernatant (cytoplasmic fraction, "Cyto") was collected and stored at -80°C. The remaining nuclear pellet was resuspended in ice-cold NER buffer supplemented with protease inhibitors, vortexed intermittently over 40 minutes, and centrifuged at 16,000 × g for 10 minutes at +4°C. The supernatant containing nuclear proteins ("Nuc") was collected and stored at -80°C until analysis. Protein concentrations were determined using the Pierce BCA Protein Assay Kit (Thermo Fisher Scientific).

#### Western Blot Analysis

Equal amounts of protein (30µg) from cytoplasmic and nuclear fractions were separated by SDS-PAGE on 4-12% gradient polyacrylamide gels (Biorad) and transferred onto nitrocellulose membranes (Biorad). Membranes were blocked in 5% non-fat dry milk in Tris-buffered saline containing 0.1% Tween-20 (TBS-T) for 1 hour at room temperature. Membranes were incubated overnight at +4°C with primary antibody against c-MAF (1:1000, rabbit polyclonal, A300-613A, Thermo Fisher Scientific/Bethyl Laboratories). Following three washes in TBS-T, membranes were incubated with HRP-conjugated anti-rabbit secondary antibody (1:25000, SantaCruz) for 1 hour at room temperature. Protein bands were visualized using enhanced chemiluminescence (ClarityECL, Biorad) and imaged using a Fusion imaging system (Vilber-Lourmat).

#### Immunofluorescence

Isolated fibers and cryosections were processed for immunofluorescence using substrate-specific protocols. Isolated fibers were blocked and permeabilized in blocking buffer containing 0.5% Triton X-100, 0.1 M glycine, and 5% normal goat serum (NGS) in PBS for 30 minutes at room temperature. For cryosections, sections were first brought to room temperature and rehydrated in PBS for 10 minutes. For c-MAF immunostaining on cryosections, slides were briefly fixed by immersion in 4% PFA in PBS for 10 seconds at room temperature immediately after thawing, then washed three times in PBS. This brief fixation step was critical for optimal c-MAF detection, as prolonged fixation reduced nuclear signal intensity while absence of fixation resulted in loss of antibody specificity. Cryosections were then blocked and permeabilized in blocking buffer containing 0.2% Triton X-100 and 5% NGS in PBS for 30 minutes at room temperature.

Both isolated fibers and cryosections were incubated with primary antibodies overnight at room temperature. For isolated fibers, antibodies included c-MAF (1:100, HPA028289, Sigma-Aldrich), FOXO3A (1:400, 75D8, Cell Signaling), α-Tubulin (1:500, GT114, GeneTex), Tubulin-β3 (1:250, Tuj1, BioLegend), acetylated-Tubulin (1:500, T6793, Sigma), Tubulin-Tyrosinated (1:500, YL 1/2 RAT, Sigma), Neurofilament-M (1:200, Poly28410, BioLegend), Synaptophysin (1:500, 7H12, Cell Signaling), detyrosinated-Tubulin (1:200, ab3201, Sigma), and c-FOS (1:200, ab208942, Abcam). For cryosections, antibodies included MYH2 (1:100, SC-71 clone, DSHB), MYH4 (1:200, BF-F3 clone, DSHB), MYH1 (1:100, 6H1 clone, DSHB), MYH7 (1:40, BA-F8, DSHB), Laminin (1:750, L9393, Sigma), FOXO3A (1:400, 75D8, Cell Signaling), and c-MAF (1:100, HPA028289, Sigma-Aldrich). Since MYH1 and MYH4 antibodies are of the same isotype (IgM), they were detected on successive serial sections.

Following three washes in PBS, samples were incubated with appropriate Alexa Fluor-conjugated secondary antibodies (1:1000, Thermofisher) for 2 hours at room temperature. Nuclei were counterstained with DAPI or Hoechst (1:10000). For neuromuscular junction visualization, α-bungarotoxin coupled with fluorophore 488 or 555 nm was added (1:1000, Invitrogen). For cryosections, WGA (1:250) was also included during secondary antibody incubation. Following three final washes in PBS, isolated fibers were additionally washed three times in blocking buffer (5% NGS, 0.1% Triton X-100 in PBS) for AChR turnover experiments. All samples were mounted on glass slides in Mowiol mounting medium.

Images were acquired using substrate-appropriate microscopy systems. For isolated muscle fibers, images were obtained using a spinning disk confocal microscope (IXplore model, Olympus manufacturer) equipped with a 60× objective. Z-stacks were collected at 0.27 µm intervals across the entire fiber depth. For cryosections, images were acquired using a widefield fluorescence microscope (BX63 model, Olympus manufacturer) equipped with a 10× objective. Whole muscle sections were captured using tile scanning and stitching. For AChR turnover experiments, images were acquired using a confocal microscope with a 60× objective. A minimum of 30 NMJ were analyzed per condition. All images were processed using FIJI software.

#### RNA In Situ Hybridization

RNA fluorescence in situ hybridization was performed on isolated muscle fibers using the RNAscope Multiplex Fluorescent Reagent Kit v2 (Advanced Cell Diagnostics, Bio-Techne) according to the manufacturer’s instructions for adherent cells. Briefly, isolated fibers were mounted on Superfrost Plus glass slides and air-dried. Fibers were dehydrated through an ethanol series (50%, 70%, 100%), and air-dried. Fibers were treated with Protease III for 40 minutes at +40°C, then hybridized with target probes for *Myh4* (pre-mRNA, 539391-C3 - Mm-Myh4-O2-C3), *Myh1* (pre-mRNA, 539381 - Mm-Myh1-O2), *Myh2* (pre-mRNA, 539031-C2 - Mm-Myh2-O1-C2), *Etv5* (mRNA, 316961-C3 - Mm-Etv5-C3) and *c-Fos* (mRNA, 316921 - Mm-Fos) for 2 hours at +40°C in a HybEZ oven. Following hybridization, signal amplification was performed using sequential amplification steps (Amp1, Amp2, Amp3) according to the manufacturer’s protocol. Fluorescent signals were developed using TSA Plus fluorophores. Nuclei were counterstained with DAPI. Slides were mounted with Mowiol mounting medium.

### In Vivo Experiments

#### Animal Models and Tamoxifen Administration

*c-Maf^flox/flox^;HSA-Cre*, *c-Maf^flox/flox^;HSA-Cre*ERT2 (Wende et al., 2012), *EnhA^-/-^* mutant mice (Dos Santos et al., 2022) and their littermate controls were maintained on a C57Bl6/N genetic background and housed in EOPS conditions at the Cochin Institute: they were maintained at a temperature of 22+/−2 °C, with 30 to 70% humidity and with a dark/light cycle of 12 h/12 h. The mice on a reversed cycle were kept for at least three weeks on a 12h/12h light/dark cycle in a specific room before tissue sampling. For Tamoxifen inducible *c-Maf^flox/flox^;HSA-Cre*ERT2 mice were administered tamoxifen (20 mg/kg body weight, Sigma-Aldrich) by intraperitoneal injection for five consecutive days. Muscles were collected at 7 days, 15 days, and 1-month post-injection. SOD1-G93A ALS mice (B6SJL-Tg(SOD1*G93A)1Gur/J, Jax No: 002726) were obtained from The Jackson Laboratory.

#### *In Vivo* Labeling of Acetylcholine Receptors

To assess acetylcholine receptor (AChR) turnover at neuromuscular junctions, a dual α-bungarotoxin (α-BTX) labeling protocol was employed. For combined denervation experiments, the initial α-BTX injection was performed 5 days post-Tamoxifen injection. Tibialis anterior (TA) muscles were injected with 25 pmol of fluorescently conjugated α-bungarotoxin (α-BTX-Alexa Fluor 488, Thermo Fisher Scientific) in 20 µl of PBS (final concentration 10 µg/ml, diluted 1:100 from stock). Ten days later, muscles were injected with 25 pmol of α-BTX conjugated to a different fluorophore (α-BTX-Alexa Fluor 555, Thermo Fisher Scientific) in 20 µl of PBS. One hour following the second injection, mice were euthanized.

#### Electrical Stimulation Protocol

Experiments were conducted on C57BL/6NCrl males (Janvier Labs, Le Genest-Saint-Isle, France) at 9 weeks of age. Mice were housed in an environment-controlled facility (12-12 hour light-dark cycle), received water and standard food ad libitum. All of the experiments and procedures were conducted in accordance with the guidelines of the local animal ethics committee of the University Claude Bernard Lyon 1 and in accordance with French and European legislation on animal experimentation and approved by the ethics committee CEEA-55 and the French ministry of research (APAFIS#22109).

Mice were initially anesthetized in an induction chamber using 4% isoflurane. The right hindlimb was shaved before an electrode cream was applied over the plantar flexor muscles to optimize electrical stimulation. Each anesthetized mouse was placed in a supine position in a noninvasive ergometer (NIMPHEA_Research, All Biomedical SAS, Grenoble, France). Throughout a typical experiment, anesthesia was maintained by air inhalation through a facemask continuously supplied with 1.5-2.5% isoflurane. The right hindlimb was stabilized and placed on two surface electrodes: one under the knee and one on Achille’s tendon to activate the plantar flexors. The right foot was firmly immobilized through a rigid slipper on a pedal connected to a torque sensor allowing for plantar flexor torque measurement. The foot was set at an angle of 15° of plantarflexion. The right knee was also firmly maintained using a rigid fixation in order to optimize maximal isometric torque recordings.

Transcutaneous single twitch stimulation was first elicited on the plantar flexor muscles using a constant-current stimulator (Digitimer DS7AH, Hertfordshire, UK; maximal voltage: 400 V; 0.2 ms duration, monophasic rectangular pulses). The individual maximal current intensity was identified by progressively increasing the stimulation until no further increase in peak twitch torque was detected. This intensity was then maintained to assess maximal isometric torque in response to a 250-ms, 100 Hz tetanic stimulation train. Following this assessment, mice were exposed to one of two stimulation protocols: a low-frequency protocol (LF; 20 Hz, 10 s on/20 s off, 30 min) or a high-frequency protocol (HF; 100 Hz, 0.6 s on/59.4 s off, 30 min). Torque data was sampled at 1000 Hz with a PowerLab8/35 (ADinstruments, Sydney, Australia), analyzed with LabChart software (v8.1.17 ADInstruments, Sydney, Australia) and normalized to maximal torque values.

### Genomics and Transcriptomics

#### RNA Extraction and Quality Control

Total RNA was extracted from frozen TA muscle tissue using TRIzol reagent (Invitrogen). Tissue samples were lysed in TRIzol using a TissueLyser (Qiagen) for mechanical homogenization, followed by phase separation with chloroform. RNA was precipitated with isopropanol, and the resulting pellet was washed three times with 75% ethanol to ensure removal of contaminants. The RNA pellet was air-dried and resuspended in nuclease-free water. RNA concentration was determined using a NanoDrop spectrophotometer, and RNA integrity was assessed using an Agilent 2100 Bioanalyzer. Only samples with RNA Integrity Numbers (RIN) ≥7.0 were used for library preparation.

#### Reverse Transcription and Quantitative PCR

500 ng of RNA was reverse transcribed into complementary DNA (cDNA) using the SuperScript reverse transcription kit (Invitrogen) according to the manufacturer’s instructions. Quantitative PCR (qPCR) was performed using SYBR Green I Master Mix (Roche) on a LightCycler real-time PCR system. Melting curve analysis was performed after each run to verify amplicon specificity and absence of primer dimers. Gene expression levels were normalized to the housekeeping gene 36B4 (Rplp0, ribosomal protein lateral stalk subunit P0) and calculated using the 2^(-ΔΔCt) method.

#### Bulk RNA Sequencing and Analysis

Fastq files were aligned using STAR algorithm (version 2.7.6a), on the Ensembl release 101 reference. Reads were then counted using RSEM (v1.3.1) and the statistical analyses on the read counts were performed with R (version 3.6.3) and the DESeq2 package (DESeq2_1.26.0) to determine the proportion of differentially expressed genes between two conditions. We used the standard DESeq2 normalization method (DESeq2’s median of ratios with the DESeq function), with a pre-filter of reads and genes (reads uniquely mapped on the genome, or up to 10 different loci with a count adjustment, and genes with at least 10 reads in at least 3 different samples). Following the package recommendations, we used the Wald test with the contrast function and the Benjamini-Hochberg FDR control procedure to identify the differentially expressed genes. R scripts and parameters are available on the platform, https://github.com/GENOM-IC-Cochin/RNA-Seq_analysis.

#### Single-Nucleus Multiome ATAC and RNA Sequencing

To comprehensively profile transcriptomic and chromatin accessibility changes following muscle denervation and tenotomy, we performed single-nucleus RNA sequencing (snRNA-seq) on *plantaris* muscle and single-nucleus multiome ATAC+RNA sequencing on tibialis anterior (TA) muscle. For denervation experiments, sciatic nerve transection was performed on 8-week-old male and female mice, and muscles were collected 7 days post-surgery. For tenotomy experiments, *plantaris* tendons were surgically severed, and muscles were collected 7 days post-surgery. Age-matched control mice were used for comparison. Muscles were rapidly dissected, snap-frozen in liquid nitrogen, and stored at -80°C until nuclei isolation. Nuclei were isolated from frozen muscle tissue using a Dounce homogenizer in ice-cold lysis buffer (10mM Tris-HCl pH7.4;10mM NaCl;3mM MgCl2;0.1% NP40;1mM DTT,40u/uL RNAse inhibitor), followed by two filtering through 70 μm and 40 μm cell strainers to remove debris and myofiber fragments and centrifuged 10min at 500g. The pellet is washed in PBS;1% BSA;1 u/uL RNAse inhibitor, centrifuged and resuspended in wash buffer. After incubation with 7AAD, 800000 nuclei were sort on an Aria and centrifuged to perform permeabilization and transposition according 10X genomics instructions.

Single-nucleus Multiome ATAC and RNA libraries were prepared using the 10x Genomics Chromium Single Cell Multiome ATAC + Gene Expression platform according to the manufacturer’s protocol. Briefly, approximately 10,000 nuclei per sample were loaded onto the Chromium Controller to generate gel bead-in-emulsions (GEMs). Following GEM generation, nuclei were barcoded and ATAC and RNA libraries were prepared in parallel. ATAC libraries were constructed using tagmentation-based methods to capture accessible chromatin regions, while RNA libraries captured nascent transcripts using polyA selection. Libraries were sequenced on an Illumina NovaSeq 6000 platform targeting 20,000 read pairs per nucleus for RNA and 25,000 read pairs per nucleus for ATAC.

Raw sequencing data were processed using Cell Ranger ARC (10x Genomics) to align reads to the mouse reference genome (mm10) and quantify gene expression and chromatin accessibility simultaneously. For RNA data, reads were aligned using STAR and gene counts were quantified using the mm10 Ensembl v79 annotation. For ATAC data, reads were aligned using BWA-MEM and peaks were called using MACS2. Filtered feature-barcode matrices were generated for downstream analysis. Nuclei were excluded if they met any of the following criteria: fewer than 200 or greater than 2,500 RNA features detected, mitochondrial transcript percentage greater than 5%, fewer than 100 or greater than 10,000 ATAC peaks detected, TSS enrichment score less than 2, or nucleosome signal greater than 10. TSS enrichment and nucleosome signal were calculated using the Signac package to ensure high-quality ATAC data.

RNA data were analyzed using Seurat v4. Normalization was performed using SCTransform with regression of RNA feature count and mitochondrial percentage to remove technical variation. Variable features (n=3,000) were identified, and principal component analysis (PCA) was performed on the scaled data matrix. To integrate data across experimental conditions (control, denervation, tenotomy) and muscle types (*plantaris*, TA), we applied Harmony batch correction using the first 25 principal components. Harmony iteratively corrected batch effects by identifying and removing batch-specific variation while preserving biological signals. UMAP dimensionality reduction was performed on the Harmony-corrected embeddings for visualization. ATAC data were processed using the Signac package. Following quality control, accessible chromatin regions were merged across all samples to create a unified peak set. Peaks were filtered to retain only those on standard chromosomes and with widths between 20 and 10,000 bp. Term Frequency-Inverse Document Frequency (TF-IDF) normalization was applied, where term frequency represents the number of fragments in a peak per nucleus normalized by total fragment counts, and inverse document frequency reflects the proportion of nuclei with any fragment in the peak. Singular value decomposition (SVD, also called Latent Semantic Indexing) was performed for dimensionality reduction, excluding the first component which typically correlates with sequencing depth. Components 2-20 were selected based on elbow plots and depth correlation analysis. Harmony integration was applied to the LSI embeddings using dimensions 2-20 to correct for batch effects across experimental conditions.

To jointly analyze RNA and ATAC modalities, we applied weighted nearest neighbor (WNN) analysis using Seurat’s FindMultiModalNeighbors function. This approach constructs a unified cell-cell similarity graph by calculating modality-specific weights for each nucleus based on the consistency of its nearest neighbors across RNA and ATAC data. WNN graphs were constructed using all Harmony-corrected RNA dimensions and ATAC dimensions, with modality weights stored as RNA.weight and ATAC.weight metadata. UMAP embedding and graph-based clustering were performed on the WNN graph using the Louvain algorithm at resolution 0.8. Cluster resolution was optimized using clustree analysis to identify stable cluster assignments across multiple resolution parameters.

Fiber type-specific differential expression analysis was performed to identify genes dysregulated in denervated versus control myonuclei within each fiber type (IIa, IIx, IIb). For each comparison, we used the Wilcoxon rank-sum test implemented in the presto package for computational efficiency. Genes were considered differentially expressed if they exhibited |log2FC| > 0.25, adjusted p-value < 0.01 (Bonferroni correction), area under the ROC curve (AUC) > 0.65, and were detected in at least 25% of nuclei in either comparison group. To identify fiber type-specific denervation signatures, we compared differentially expressed gene lists between denervation (sciatic nerve transection) and tenotomy conditions. Genes that were significantly upregulated or downregulated specifically in denervated IIb nuclei relative to both control and tenotomized conditions were classified as denervation-specific IIb genes. To identify master regulatory factors driving fiber type-specific responses to denervation, we performed upstream regulator (UR) analysis using Ingenuity Pathway Analysis (IPA, QIAGEN). Differentially expressed gene lists from IIa, IIx, and IIb myonuclei were uploaded separately along with their fold change and adjusted p-values. IPA’s Upstream Regulator Analysis algorithm predicts transcription factors, signaling molecules, and other regulators that can explain the observed gene expression changes based on compiled literature-derived regulatory relationships. Z-scores were calculated to assess the activation state of each predicted upstream regulator, with |Z-score| > 2 indicating significant activation or inhibition. To identify upstream regulators specific to denervation (versus tenotomy) and specific to type IIb nuclei (versus IIa and IIx), we compared UR lists across conditions and fiber types. URs present in denervated IIb nuclei but absent in tenotomized muscle and absent in denervated IIa and IIx nuclei were classified as IIb-specific denervation upstream regulators.

#### ChIP-seq Analysis

ChIP-seq peak coordinates for c-MAF (skeletal muscle and Th17 cells), glucocorticoid receptor (GR, muscle), and pioneer transcription factors (SOX2, KLF4, ETV5) were extended ±100 bp from peak summits. Genomic sequences were extracted from the mm10 reference genome using BSgenome.Mmusculus.UCSC.mm10. Transcription factor binding motifs were identified using the Biostrings matchPattern function with strand-specific scanning, allowing one mismatch per motif to account for natural binding site variation. TF-specific consensus sequences were: c-MAF (TGCTGAC, half-MARE), GR (AGAACA, half-GRE), ETV5 (ACCGGAAGT, ETS core), SOX2 (CATTGT, SOX core), and KLF4 (CCGCCC, KLF core).

For each ChIP-seq peak containing multiple motif occurrences, pairwise spacing distances were computed as the genomic distance between the 3’ end of the upstream motif and the 5’ start of the downstream motif, with a maximum distance threshold of 200 bp. Spacing distributions were binned at 5 bp resolution and analyzed across three biologically relevant nucleosome-compatible spacing ranges: same-gyre positioning (35-45 bp, representing motifs on the same face of the DNA helix), cross-gyre positioning (70-85 bp, representing motifs on opposite helical faces separated by ∼2 helical turns), and full-nucleosome positioning (140-155 bp, representing motifs separated by one complete nucleosomal DNA wrap). To establish null distributions for spacing frequencies, motif positions within each ChIP-seq peak were randomly shuffled 500 times while preserving the total number and genomic bounds of motifs per peak. For each spacing category, fold enrichment was calculated as the ratio of observed to median shuffled spacing frequency. Statistical significance was determined by comparing observed frequencies to the shuffled null distribution, with Bonferroni correction applied for multiple hypothesis testing across three spacing categories (α = 0.05/3 = 0.017). This approach controls for peak-specific sequence composition and motif density effects.

To quantify cooperative binding between c-MAF response elements (MAREs) and co-regulatory transcription factors, we analyzed the spatial proximity of TF binding sites (SIX1, KLF4, SOX6, TBX15, NR4A1, GR) to half-MARE motifs (TGCTGAC) within ChIP-seq peaks. For each TF-MARE pair, proximity was assessed across multiple distance thresholds (ranging from 10 to 100 bp) by counting the number of MARE sites with at least one TF binding site within the specified threshold distance. MARE-MARE pairs were defined as two half-MARE motifs occurring within 200 bp, representing potential full or composite MAF response elements. For each distance threshold, fold enrichment was calculated as the observed frequency of TF-proximal MARE pairs divided by the expected frequency under random spatial distribution. Expected frequencies were computed by randomly shuffling TF binding site positions within peak boundaries while maintaining motif counts. Statistical significance was assessed using Fisher’s exact test, comparing the observed contingency table (MARE pairs with/without proximal TF sites) against the null hypothesis of random spatial distribution. The optimal proximity threshold for each TF was defined as the distance yielding maximum fold enrichment. Cooperativity profiles were visualized as fold enrichment curves across distance thresholds, with optimal points highlighted for each TF. Multi-panel layouts enabled direct comparison of cooperativity strength and optimal binding distances across TF families. Overlay plots superimposed all TF enrichment profiles to identify shared or divergent spatial relationships with MARE sites. Summary statistics included optimal threshold distances, maximum fold enrichment values, total TF binding sites analyzed, and Fisher’s exact test p-values at optimal distances.

Genes associated with half-MARE pairs (TGCTGAC motifs occurring within 200 bp) were identified using ChIPseeker peak annotation with the TxDb.Mmusculus.UCSC.mm10.knownGene database (TSS region: -3000 to +3000 bp). Half-MARE pairs were stratified into three nucleosome-compatible spacing categories: same-gyre positioning (35-45 bp, motifs on the same DNA helical face), cross-gyre positioning (70-85 bp, motifs on opposite helical faces separated by approximately 2 helical turns), and full-nucleosome positioning (140-155 bp, motifs separated by one complete nucleosomal DNA wrap of 147 bp). To resolve genes associated with multiple spacing categories due to overlapping MARE pairs within a single peak, a priority assignment system was implemented: cross-gyre > full-nucleosome > same-gyre. This hierarchy assigns each gene to its most prominent spacing category, creating mutually exclusive gene sets for downstream enrichment analysis.

Gene Ontology enrichment analysis was performed using clusterProfiler with Biological Process, Cellular Component, and Molecular Function terms from org.Mm.eg.db. For each significantly enriched GO term (adjusted p < 0.05, FDR correction), we calculated background-adjusted enrichment scores rather than assuming uniform distribution across spacing categories. The background distribution was empirically determined from the observed gene counts in each spacing category (same-gyre, cross-gyre, full-nucleosome), and enrichment was calculated as the ratio of observed to expected gene counts based on this background. For each GO term, Fisher’s exact tests were performed independently for each spacing category to test enrichment against the combined background of the other two categories, providing spacing-specific significance values. Chi-square tests with background-adjusted expected proportions were additionally performed for GO terms with at least 10 genes and expected counts ≥1 per category. A GO term was classified as having a dominant spacing bias if its enrichment exceeded 1.2-fold in one category and this category showed the maximum enrichment value. Enriched GO terms were organized into functional meta-categories based on biological themes: Skeletal Muscle (differentiation, development, contraction), Neuromuscular Junction (synapse organization, axon guidance, neuromuscular process), Metabolism (phosphate metabolism, energy generation, catabolic processes), Vesicle (secretion, transport, endoplasmic reticulum stress), and Signaling (MAPK, calcineurin-NFAT, intracellular pathways). To identify GO terms with differential enrichment patterns across nucleosome spacing configurations, unbiased hierarchical clustering was performed on the matrix of fold enrichment values (same-gyre, cross-gyre, full-nucleosome) using Euclidean distance and complete linkage. GO terms exhibiting pronounced spacing bias (enrichment >1.5 in one category relative to others, adjusted p < 0.05) were classified as biased terms and subjected to secondary clustering within functional modules. The top five most significantly enriched terms per functional category were selected for integrated visualization based on gene ratio (proportion of category genes in the enriched term) and absolute gene count.

---

**Supplementary Figure 1.**
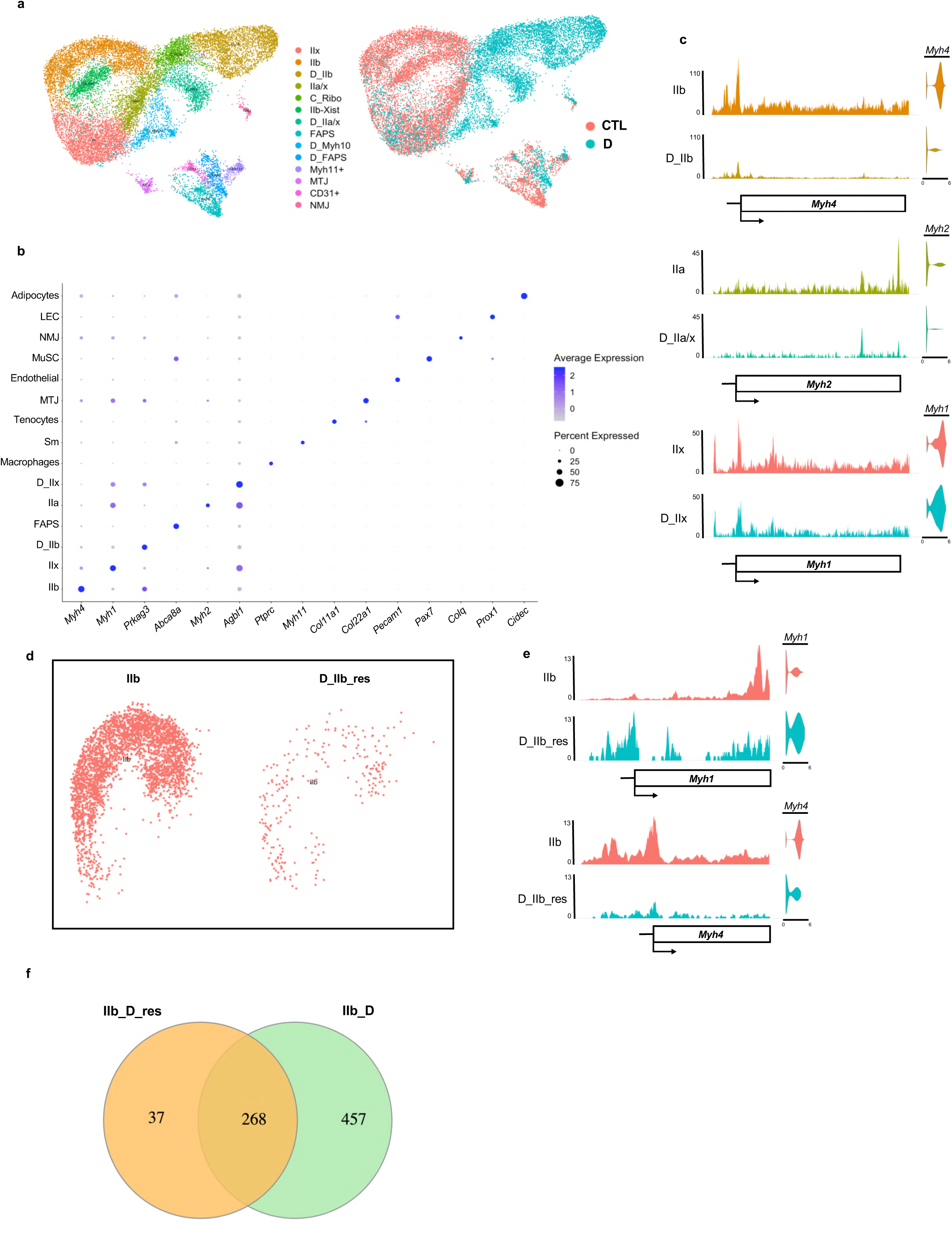
Single-nucleus Multi-omic analysis revealed resilient type IIb myonuclei population retaining transcriptional identity during denervation. **a**, weighted nearest neighbor (WNN) UMAP integrating snATAC-seq and snRNA-seq from tibialis anterior muscle 7 days post-denervation. Left panel colored by cluster identity shows major populations. Right panel colored by experimental group shows control (CTL, red) versus denervated (D, blue) **b**, Dot plot of marker gene expression across cell clusters. Y-axis lists 15 clusters; X-axis displays marker genes. Dot size indicates percent expression; color intensity (purple gradient) shows average expression level. Validates cluster identities through fiber type-specific *Myh* expression and cell type-specific markers. **c**, Coverage plots showing chromatin accessibility at f*Myh* locus for *Myh4*/IIb, *Myh2*/IIa, and *Myh1*/IIx in control (CTL) and denervated (D) TA, with at the right violin plots showing the expression level of the corresponding gene. Below, in black each gene is represented. **d**, UMAP subset focusing on type IIb myonuclei. Left (CTL) shows control IIb nuclei right (D) shows resilient denervated IIb nuclei that retain spatial clustering with control population (D_IIb_res), suggesting maintenance of transcriptional/chromatin accessibility identity despite denervation. **e**, Coverage plots comparing *Myh1* and *Myh4* loci in resilient denervated population. Cyan tracks showing denervated nuclei; red tracks showing control nuclei. Violin plots (right) indicate gene expression. *Myh1* and *Myh4* genes are presented below corresponding the coverage plots. **f**, Venn diagram comparing differentially expressed genes between IIb denervated nuclei which cluster with the control (resilient myonuclei, 305 genes, orange) and those (non-resilient, 725 genes, green) that do not cluster with the control. Overlap shows 268 shared dysregulated genes; 37 unique to resilient, 457 unique to non-resilient.

**Supplementary Figure 2.**
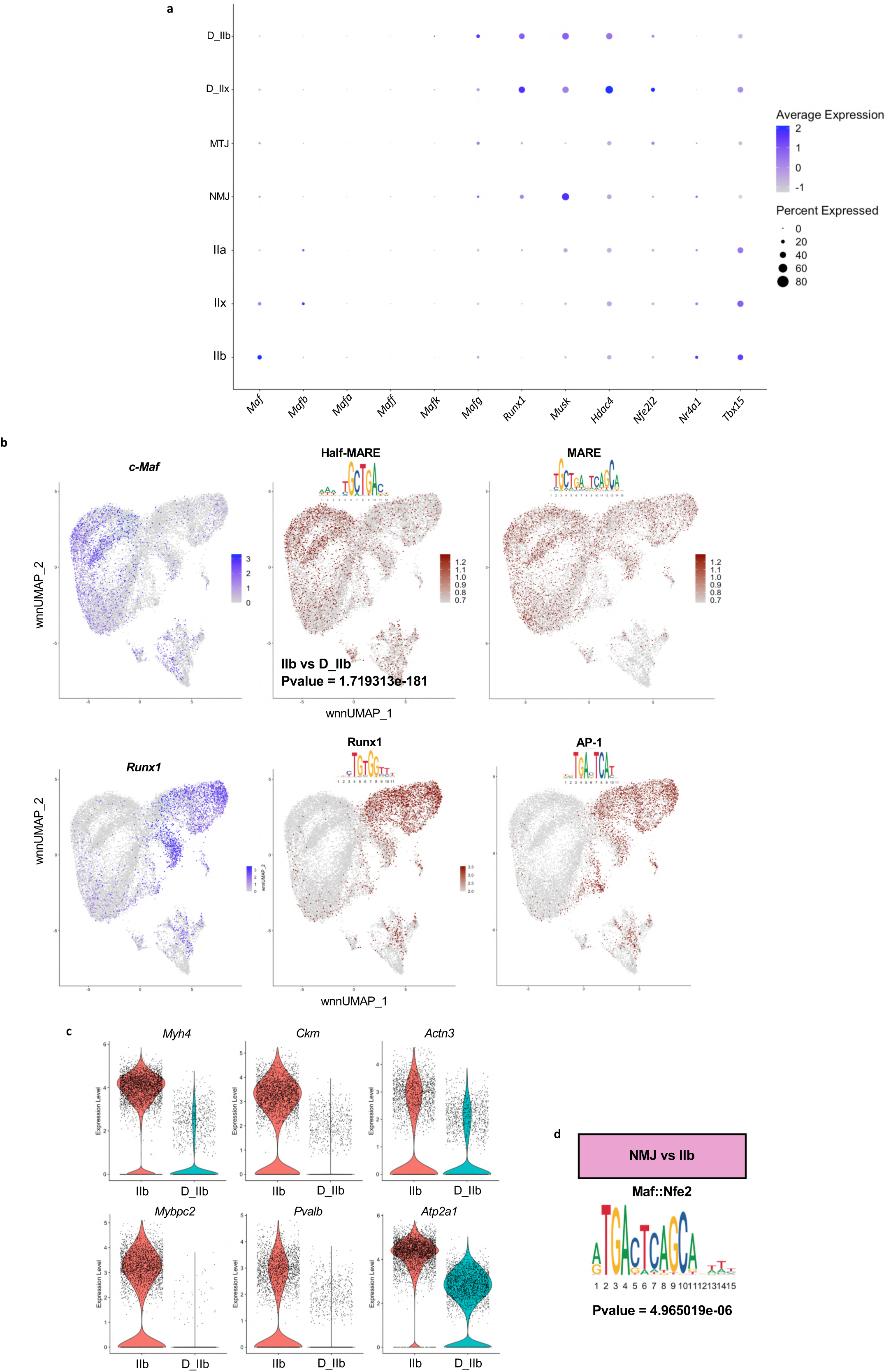
*c-Maf* expression and transcription factor motif accessibility patterns in denervated tibialis anterior muscle. **a**, Dot plot showing expression of *Maf* family transcription factor, *Runx1*, *Hdac*4, *Nr4a1*, *Tbx15*, Musk*, Nfe2l2* expression across distinct myonuclei populations. Y-axis lists seven cell populations. X-axis displays gene names. Dot size indicates percentage of cells expressing each gene (legend shows 0, 10, 20, 30, 40 percent). Dot color intensity (purple gradient) represents average expression level (scale 0 to 2 arbitrary units). Populations include control (IIa, IIx, IIb) and denervated (D_IIb, D_IIx) myonuclei plus specialized populations (NMJ, MTJ). **b**, Six UMAP feature plots displaying gene expression and chromatin accessibility patterns from WNN-integrated multiome dataset, as in Supplementary Figure 1a. Top row, left to right: *c-Maf* RNA expression (purple-to-gray gradient), half-MARE motif chromatin accessibility (brown-to-gray gradient), palindromic MARE motif accessibility (brown-to-gray gradient). Bottom row, left to right: *Runx1* RNA expression (purple-to-gray gradient), RUNX1 binding motif accessibility (brown-to-gray gradient), FOS motif accessibility (brown-to-gray gradient). Color intensity indicates expression level or accessibility score per nucleus. **c,** Violin plots showing expression of type IIb fiber-specific genes in IIb myonuclei (red) versus denervated IIb myonuclei (D_IIb, cyan) from single-nucleus RNA-seq of tibialis anterior muscle at 7 days post-denervation. Denervation induces downregulation of the fast-glycolytic contractile program, including *Myh4* (type IIb myosin heavy chain), *Actn3* (alpha-actinin-3), and *Mybpc2* (myosin binding protein C2), alongside metabolic enzymes *Ckm* (muscle creatine kinase) and *Pvalb* (Parvalbumin) and the sarcoplasmic reticulum calcium ATPase *Atp2a1* (SERCA1). Each dot represents an individual myonucleus; violin width indicates cell density distribution. **d**, Chromvar analysis of MAF::NFE2 motif enrichment in NMJ compared to IIb CTL showing significant enrichment.

**Supplementary Figure 3.**
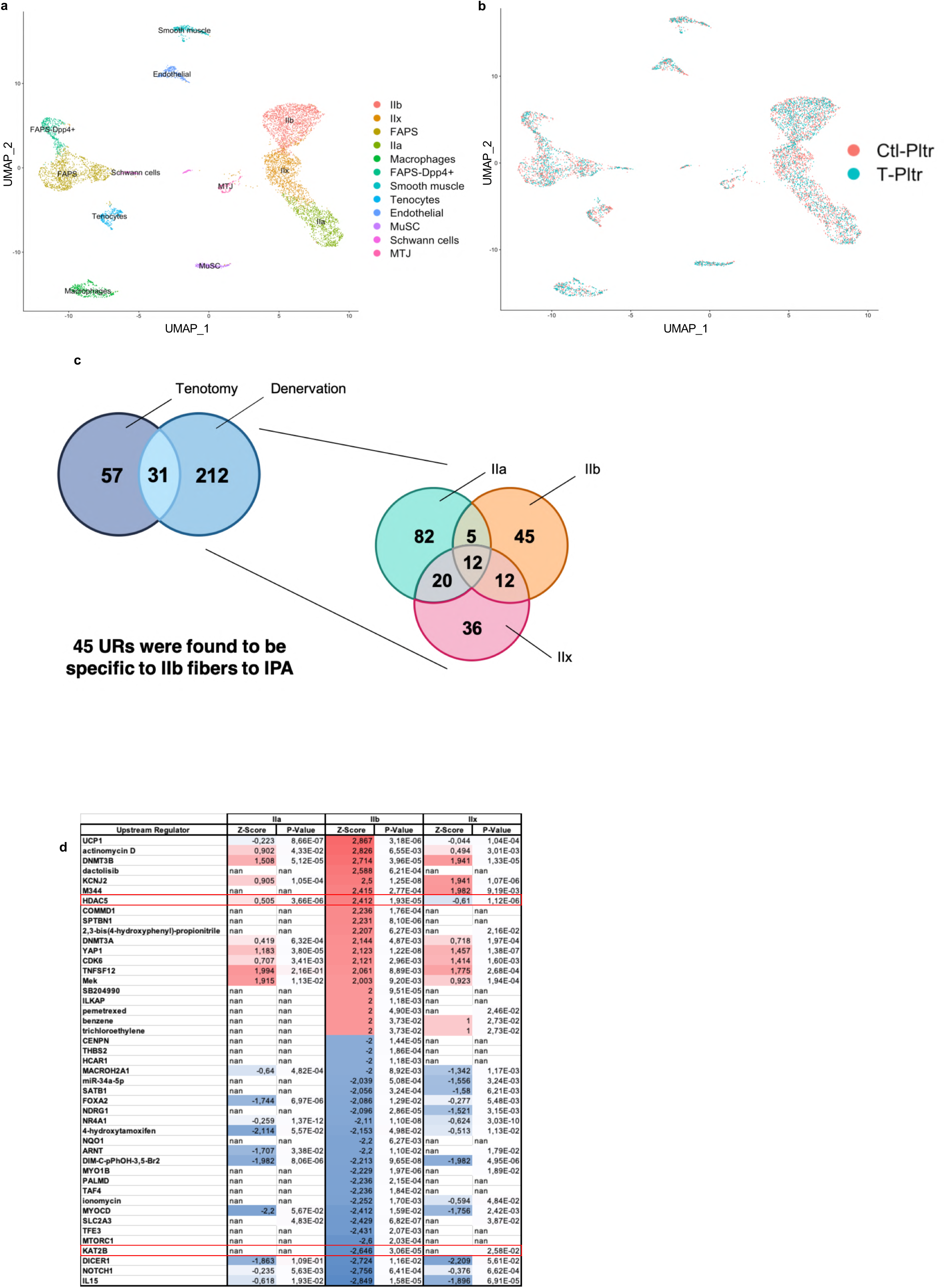
Single-nucleus RNA-seq revealed fiber type-specific transcriptional responses to mechanical unloading. **a**, UMAP visualization of single-nucleus RNA-seq from *plantaris* muscle 7 days post-tenotomy (mechanical unloading) and sham-operated controls. Cells colored by cluster identity reveal 11 distinct populations: IIb (orange, type IIb/*Myh4* myonuclei), IIx (red, type IIx/*Myh1* myonuclei), FAPs (yellow, fibro-adipogenic progenitors), IIa (green, type IIa/*Myh2* myonuclei), Macrophages (teal), FAPS-*Dpp4*+ (cyan), Neuromuscular cells (blue), Tenocytes (light blue), Endothelial (purple), MuSC (magenta, muscle stem cells), Schwann cells (pink), MTJ (pink, myotendinous junction). Myonuclei clusters segregate by *Myh* isoform expression demonstrating fiber type-specific transcriptional states. **b**, Same UMAP colored by experimental condition using group.by function. Cyan indicates Ctrl-Pltr (control *plantaris*); pink indicates T-Pltr (tenotomized *plantaris*). Both conditions distribute across all clusters, enabling paired comparison of tenotomy effects within each cell type. **c**, Venn diagram analysis of Ingenuity Pathway Analysis (IPA) upstream regulators (URs). Left: Overlap between Tenotomy (57 unique URs) and Denervation (212 unique URs) showing 31 shared regulators, indicating partially overlapping stress responses. Right: Triple Venn diagram among the 212 unique denervation URs showing fiber type-specific ones: type IIa (82 unique), IIb (45 unique), and IIx (36 unique) myonuclei. Central overlaps show shared regulators: 5 (IIa/IIb), 12 (all three), 12 (IIb/IIx), and 20 (IIa/IIx). **d**, Table of IIb-specific upstream regulators showing z-scores and p-values across fiber types (IIa, IIb, IIx). Columns display regulator name, activation z-score, and statistical significance for each subtype. Red cells indicate activated pathways; blue indicates inhibited pathways. Red rectangles highlight two key regulators: KAT2B (Lysine acetyltransferase) showing its specific inhibition in IIb fibers, and HDAC5 (class IIa histone deacetylase) showing its specific activation. This inverse regulation suggests coordinated chromatin remodeling where HDAC5 activation and KAT2B inhibition converge to alter acetylation landscapes during denervation of fast glycolytic IIb myofibers.

**Supplementary Figure 4.**
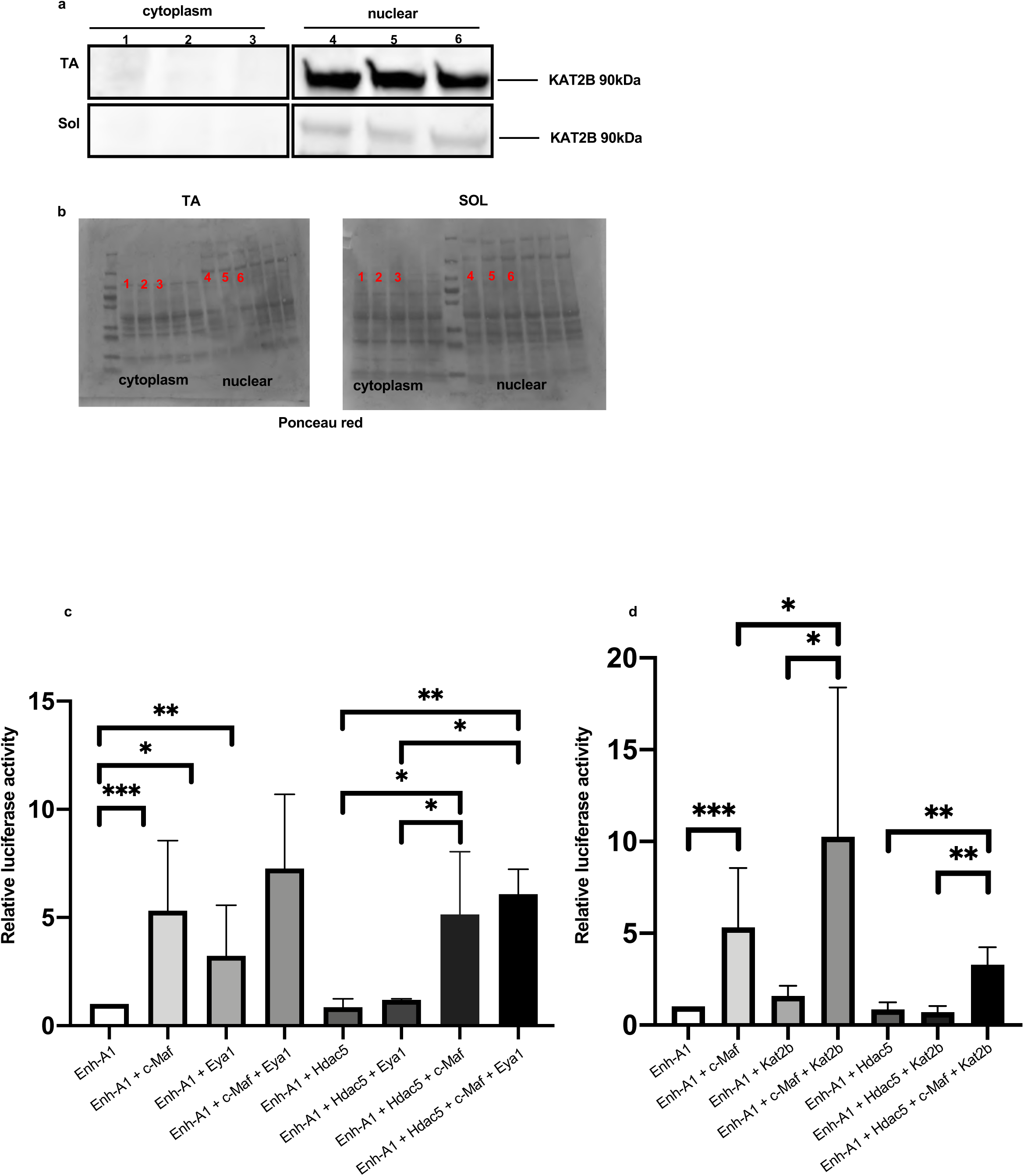
KAT2B showed fiber-type specific nuclear localization and synergizes with c-MAF to activate transcription. **a**, Western blot analysis of KAT2B (lysine acetyltransferase 2B, 100 kDa) subcellular localization in fast and slow normal muscles. Top panels: tibialis anterior (TA, fast glycolytic). Bottom panels: soleus (Sol, slow oxidative). Left columns show cytoplasmic fractions; right columns show nuclear fractions. **b**, Ponceau S total protein staining serving as loading and transfer controls for western blot membrane shown in **a,** showing protein fraction in cytoplasm and nucleus. **c**, Luciferase reporter activity of the Enh-A1 enhancer co-transfected with *c-Maf*, *Eya1*, and/or *HDac5* expression vectors. c-MAF and EYA1 activate the enhancer, while HDAC5 suppresses EYA1 induced activity but not c-MAF induced activity. **d**, Luciferase reporter activity of Enh-A1 co-transfected with *c-Maf*, *Hdac5*, and/or *Kat2b* expression vectors. KAT2b potentiates c-MAF-dependent activation but do not relieve HDAC5-mediated repression. Data are presented as mean ± s.d. relative to empty control vector. *P < 0.05, **P < 0.01, ***P < 0.001 (one-way ANOVA).

**Supplementary Figure 5.**
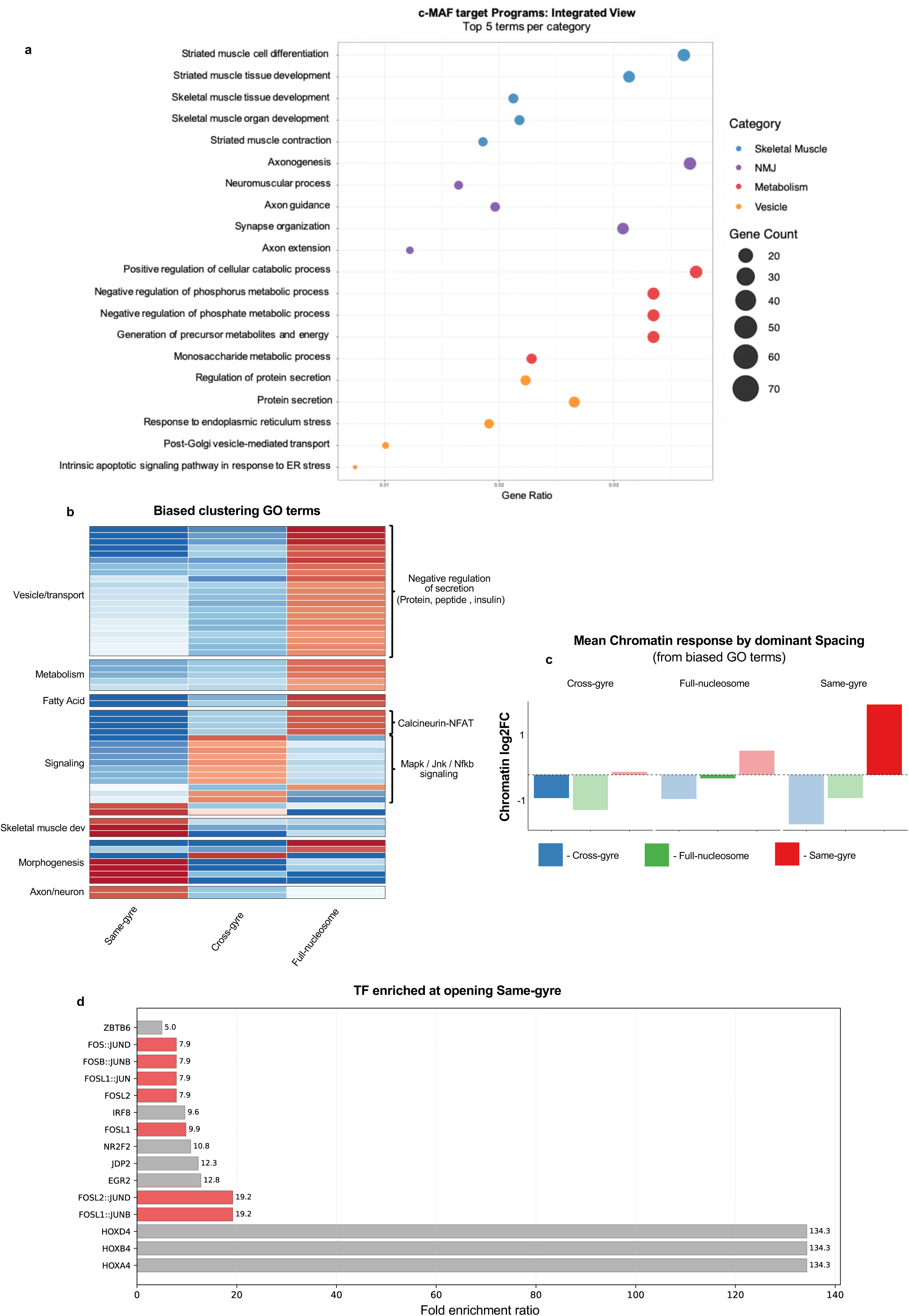
Nucleosome spacing architecture segregated c-MAF target genes by functional category and transcriptional dynamics. **a**, Dot plot showing enriched biological processes for genes associated with nucleosome compatible Half-MARE pairs (140-155 bp) from chip-seq data. Y-axis displays GO term categories ordered by significance. X-axis shows gene ratio (proportion of genes in each category). Dot size represents gene counts. Terms are displayed by four major functional modules color-coded by category: Metabolism (red), NMJ (purple), Vesicles (orange), and Skeletal muscle (blue). **b**, Heatmap of specific terms mentioned in the paper organized by pre-defined functional categories (Skeletal Muscle, Metabolism, Vesicle/Transport, Signaling, and other biological processes). Rows grouped by category rather than hierarchical clustering to highlight functional segregation. Color scale as in (e). Vesicle/Transport category (n=21 terms) shows strongest full-nucleosome enrichment (mean 1.93-fold). Signaling pathways split between cross-gyre (n=10 terms, mean 1.65-fold enrichment, activity-responsive kinase cascades) and full-nucleosome (n=5 terms, mean 2.01-fold enrichment, calcium/Wnt homeostatic signaling). Skeletal muscle differentiation terms (n=3) show exclusive same-gyre enrichment (mean 3.02-fold). **c**. Mean chromatin accessibility changes (log₂FC) in denervated IIb myonuclei for GO terms biased toward each spacing category (cross-gyre, full-nucleosome, or same-gyre). Bars represent chromatin response measured at all three spacing patterns within genes enriched for each GO term set. **d**, Transcription factor motif enrichment at same-gyre c-MAF binding sites that undergo chromatin opening upon denervation. Bars show fold enrichment ratio relative to genomic background. Red bars indicate AP-1 family heterodimers (FOS/JUN); gray bars indicate other TF families. HOX homeodomain factors show the strongest enrichment (>130-fold), followed by AP-1 heterodimers FOSL2::JUND and FOSL3::JUNB (∼19-fold).

**Supplementary Figure 6.**
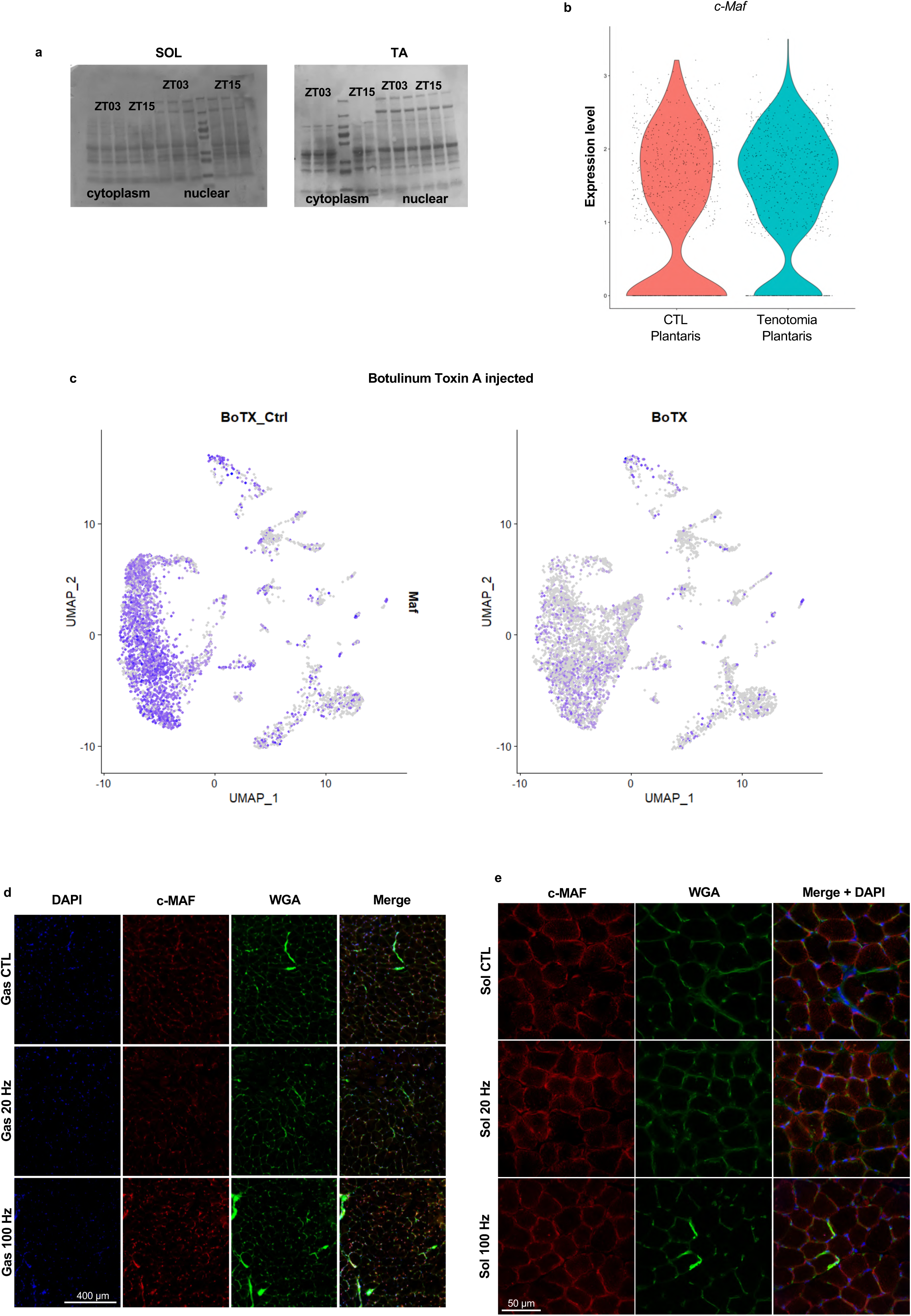
Western blot loading controls and botulinum toxin-induced c-MAF downregulation in single myonuclei. **a**, Ponceau S total protein staining serving as loading controls for western blots presented in main Figure 3 of the manuscript. Two membrane panels show protein transfer quality for soleus (SOL) and tibialis anterior (TA) muscles. Each panel displays cytoplasmic and nuclear protein fractions. **b.** Violin plots showing expression of *c-Maf* in *Myh4*+ myonuclei from single-nucleus RNA-seq in control *plantaris* and *plantaris* after tenotomy. **c**, UMAP visualization of single-nucleus RNA-seq from botulinum toxin A (BoTX)-injected muscles (Ham et al., 2025). Two Feature plots compare gene expression profiles: left panel shows BoTX_Ctrl (control-treated), right panel shows BoTX (toxin-treated). Each point represents an individual myonucleus positioned by UMAP dimensionality reduction. Color intensity (purple-to-gray gradient) indicates *c-Maf* expression level per nucleus, with "*Maf*" label on right margin. **d,** Widefield images at x10 magnification of Gastrocnemius muscle in CTL and after electrical stimulation (20 Hz and 100 Hz) stained for c-MAF (red), WGA (green) and Dapi (blue). **e**, Spinning-disk confocal images at x60 magnification of soleus muscle after electrical stimulation same protocol and staining as in d.

**Supplementary Figure 7.**
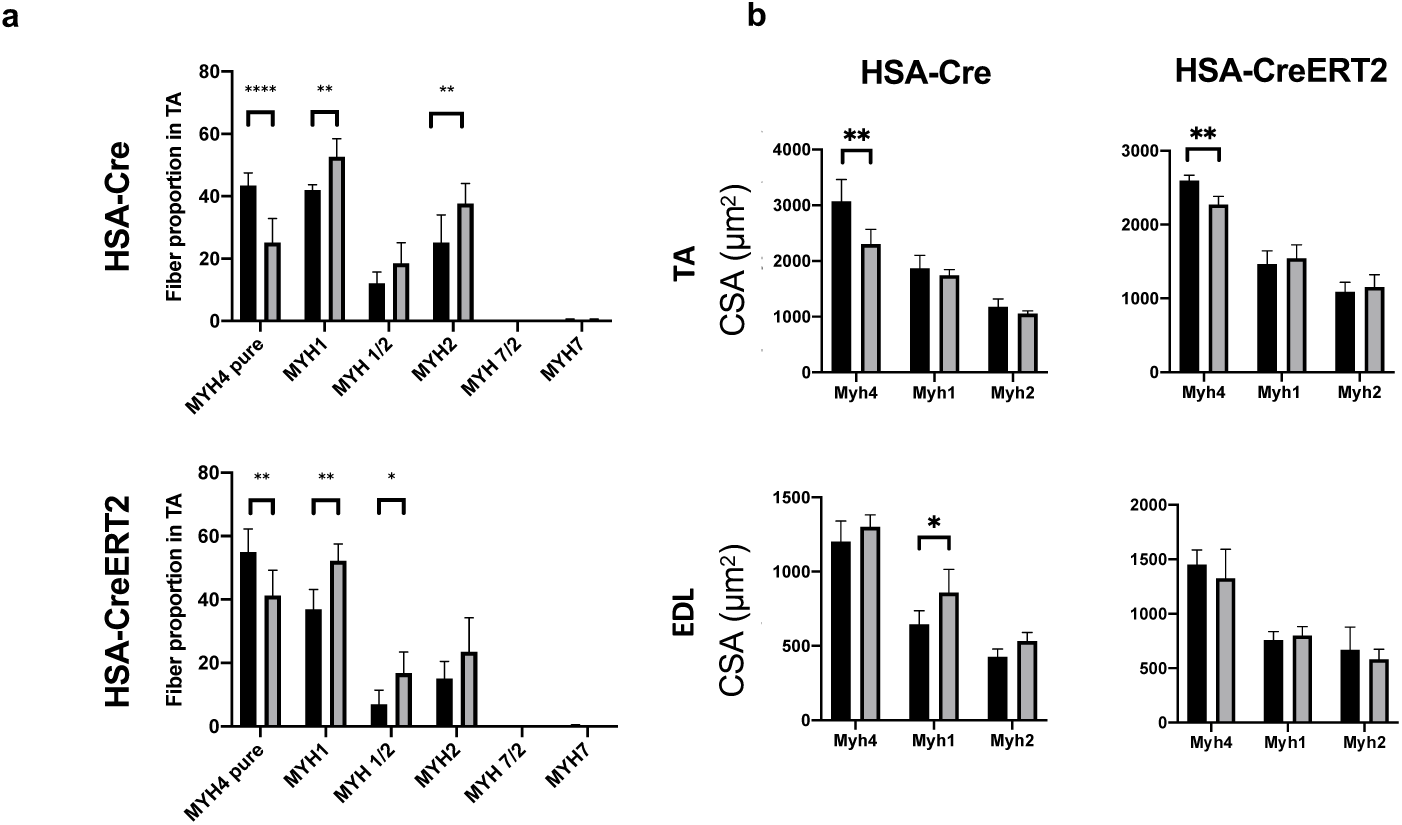
Muscle fiber type composition and cross-sectional area revealed regional vulnerability in *c-Maf* muscle-specific mutant mice. **a,** Quantification of fiber type composition in tibialis anterior (TA) muscle from *c-Maf* mutant mice generated using constitutive HSA-Cre (Top) or inducible HSA-CreERT2 (Bottom), assessed by Myosin heavy chain (MYH) isoform immunostaining. Wild-type (black bars) and c-*Maf* knockout (white/gray bars) muscles show significant reduction in type IIb (MYH4) fiber content with compensatory increases in type IIx (MYH1) and type IIa (MYH2) fibers following c-*Maf* deletion. **b,** Cross-sectional area (CSA µm^2^) measurements of individual fiber types in TA (top) and EDL (bottom) muscles from HSA-Cre (left) and HSA-CreERT2 (right) mutant mice. Selective atrophy of MYH4-expressing type IIb fibers is observed exclusively in TA muscle, with no CSA changes detected in EDL, revealing region-specific vulnerability to *c-Maf* loss despite comparable fiber type composition. Note that the CSA of MYH4 myofibers is around 3000 µm^2^ in the TA, while of 1200 µm^2^ in EDL.

**Supplementary Figure 8.**
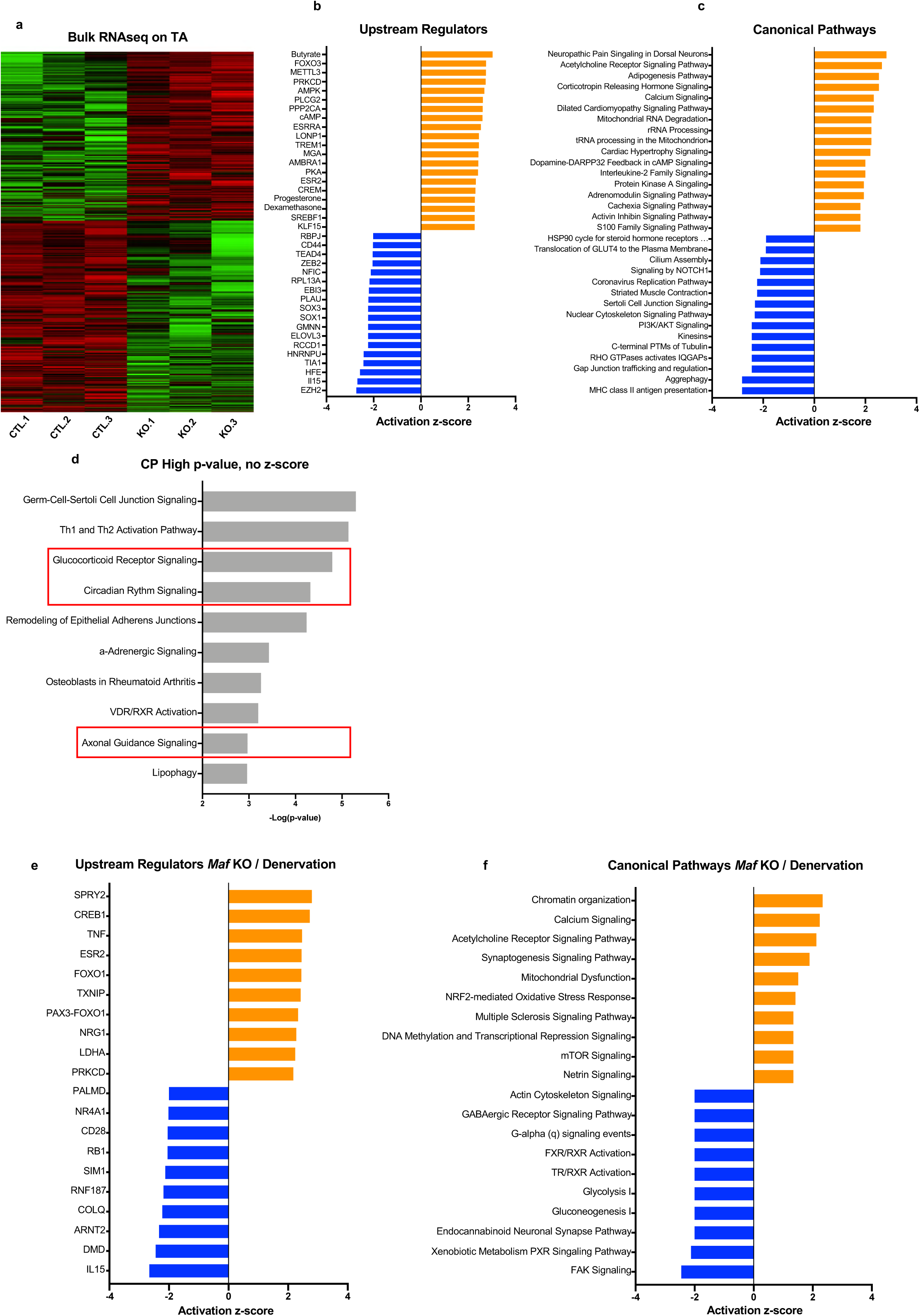
Bulk RNA-seq revealed that c-MAF regulates overlapping transcriptional programs that are downregulated during denervation and in *c-Maf* mutants. **a**, Heatmap of bulk RNA-sequencing from tibialis anterior (TA) muscle in constitutive *c-Maf* muscle-specific mutant. Rows represent differentially expressed genes hierarchically clustered by expression pattern. Columns show biological replicates. Color scale: red (low expression), green (high expression), black (intermediate). **b**, Ingenuity Pathway Analysis (IPA) upstream regulator (UR) analysis for *c-Maf* dataset. Horizontal bar graph shows activation z-scores for predicted upstream regulators. Orange bars (positive z-score) indicate activated pathways; blue bars (negative z-score) indicate inhibited pathways in mutant TA. X-axis shows activation z-score from -4 to 4. Analysis identifies transcriptional cascades dysregulated upon *c-Maf* loss. **c**, IPA canonical pathway (CP) analysis for *c-Maf* mutant showing significantly enriched pathways. Orange bars indicate activated pathways, blue bars show inhibited pathways **d**, Canonical pathways with significant p-values but no calculated z-scores. Horizontal bars show -log10(p-value) for pathways including glucocorticoid signaling, circadian rhythm signaling (red box), and axonal guidance signaling (red box). These pathways show statistical enrichment but lack directional z-scores. **e and f**, IPA comparative analysis of bulk RNA-seq during denervation and c-*Maf* mutant bulk RNA-seq. Canonical Pathway (**g**) and Upstream Regulator (**f**).

**Supplementary Figure 9.**
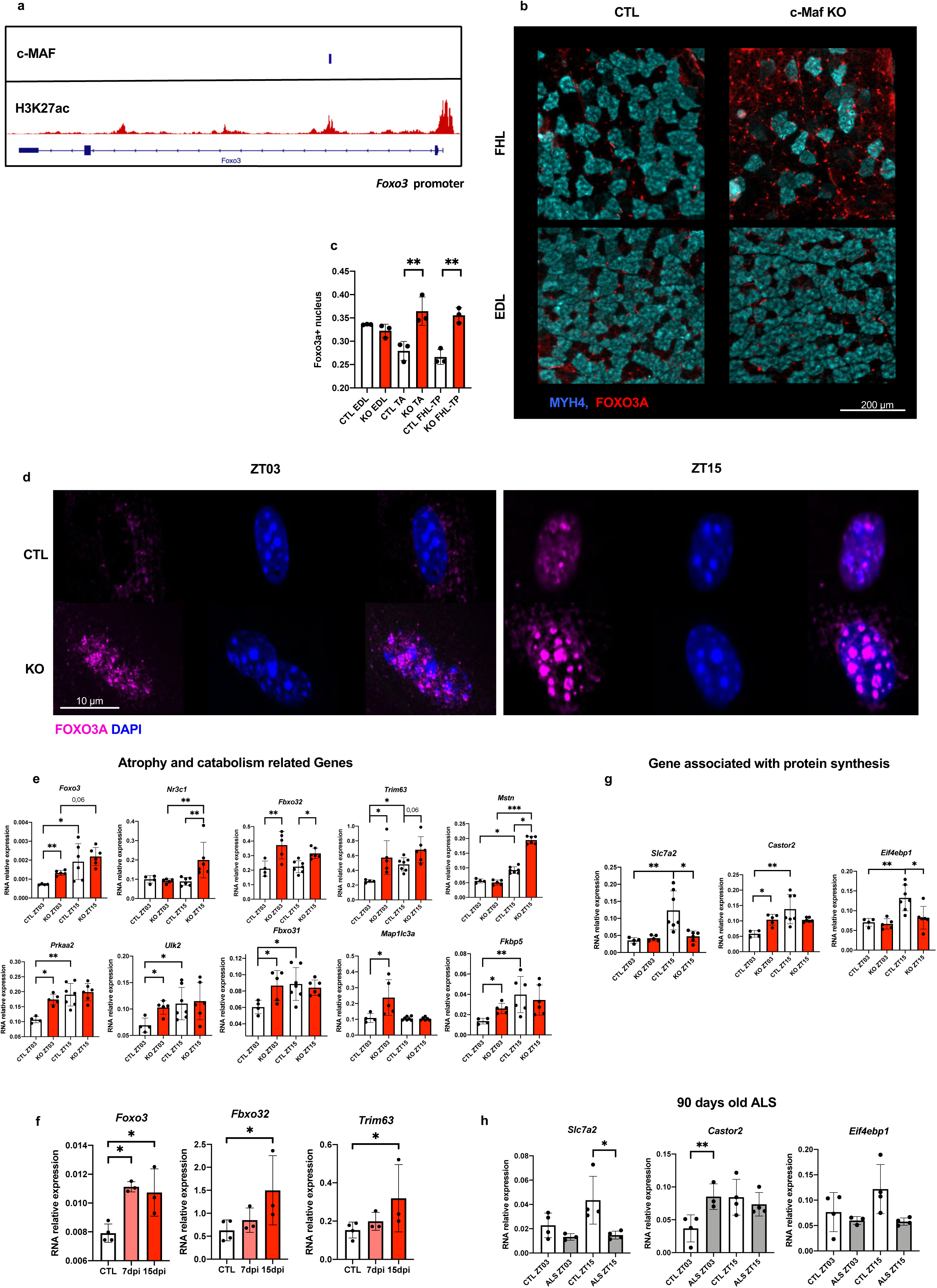
*c-Maf* loss activated FOXO3-dependent atrophy program with circadian and temporal dynamics. **a**, Chip-seq alignment of c-MAF peaks with H3K27ac histone mark on *Foxo3* gene showing c-MAF significant peak in intronic enhancer of *Foxo3.* **b**, Immunofluorescence of FOXO3 in flexor hallucis longus (FHL) and extensor digitorum longus (EDL) sections. MYH4 (cyan) and FOXO3A (red). **c**, Quantification of FOXO3-positive myonuclei across muscles (CTL-EDL, KO-EDL, CTL-TA, KO-TA, CTL-FHL-TP, KO-FHL-TP). **p < 0.01 Two-way ANOVA. **d**, Confocal imaging of isolated TA fiber nuclei showing FOXO3A (magenta) and DAPI (blue) at ZT03 and ZT15. **e**, RT-qPCR of atrophy genes *Foxo3*, *Nr3c1*, *Nr3c2*, *Fbxo32*, *Foxo1*, *Trim63*, *Fbxo31*, *Map1lc3a*, *Mstn*, *Fkbp5*) at ZT03/ZT15. White bars (CTL), red bars (KO). *p < 0.05, **p < 0.01 One-way ANOVA. **f**, RT-qPCR of protein synthesis genes (*Castor2*, *Eif4ebp1*, *Slc7a2*, *Eif4e*). **g**, Gene expression kinetics after *c-Maf* deletion. Graphs showing *Foxo3*, *Fbxo32*, *Trim63* expression at 7 and 15 dpi of tamoxifen, *p < 0.05 One-way ANOVA. **h**, Expression of *Castor2*, *Eif4ebp1* and *Slc7a2* in 90-day old CTL and ALS SOD1-G93A at ZT03/ZT15. ALS and *c-Maf* KO mice showed comparable dysregulation of those genes. *p < 0.05, **p < 0.01 One-way ANOVA.

**Supplementary Figure 10.**
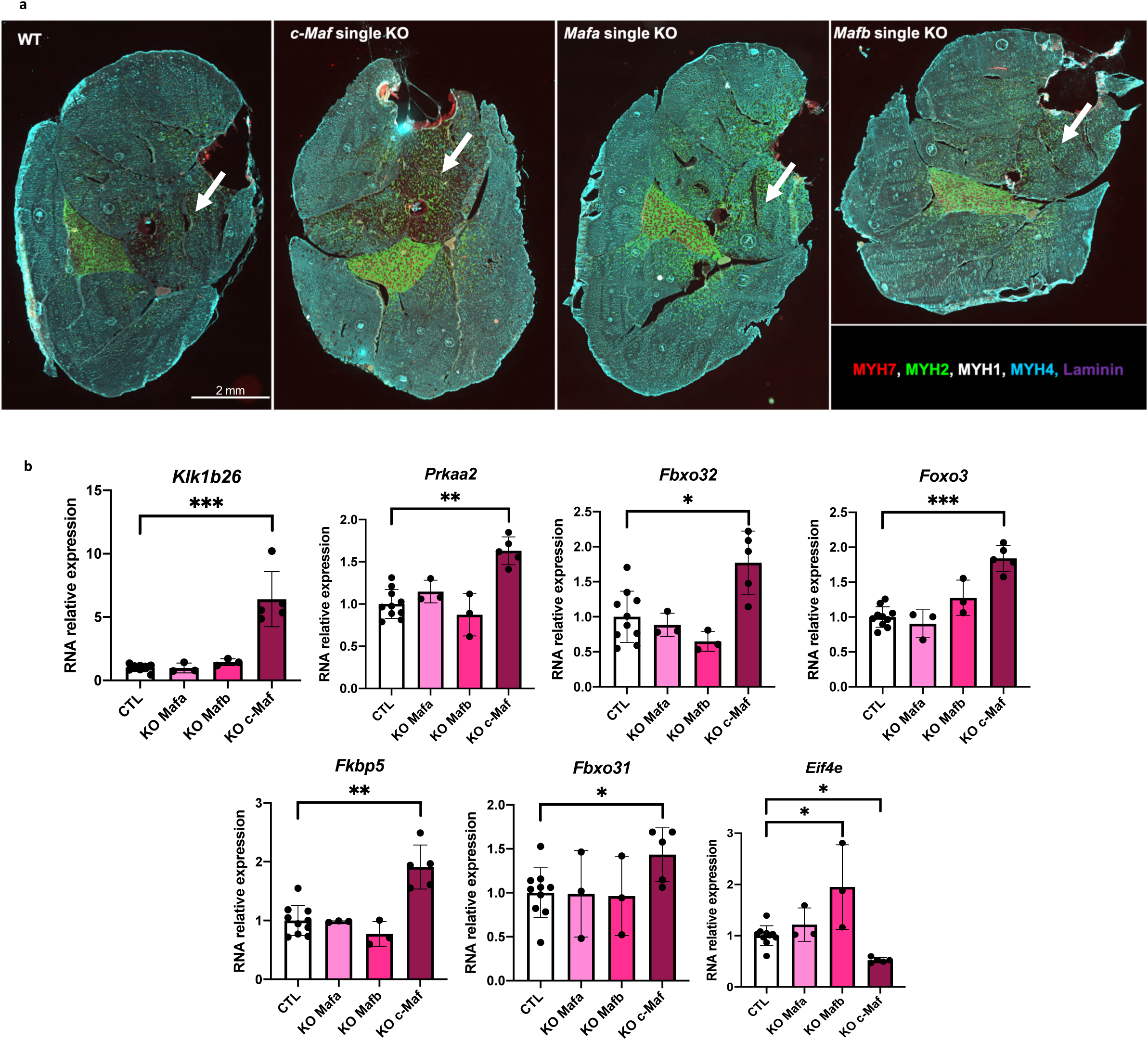
c-MAF exhibited unique muscle regulatory function distinct from MAFA and MAFB. **a**, Immunofluorescence of distal hindlimb cryosections comparing muscle-specific large *Maf* family mutant models (*HSA-Cre; c-Maf ^flox/flox^, HSA-Cre; Mafb ^flox/flox^ and Mafa* null mice). Four panels showing transverse sections from: wild-type (WT), *c-Maf* single mutant, *Mafa* single mutant, and *Mafb* single mutant mice. Myosin heavy chain (MYH) isoforms visualized by immunostaining: MYH7 (blue, slow oxidative), MYH2 (red, fast oxidative), MYH1 (white, fast oxidative-glycolytic), MYH4 (green, fast glycolytic), and Laminin (magenta, basal lamina marking fiber boundaries). White arrows indicating flexor hallucis longus (FHL) and tibialis posterior (TP) muscles showing region-specific fiber type transition in *c-Maf* mutant. **b**, RT-qPCR quantification of genes specifically dysregulated in *c-Maf* mutant compared to *Mafb* and *Mafa* mutants. Seven bar graphs show relative mRNA expression for: *Klk1b26*, *Prkaa2*, *Fbxo32*, *Foxo3*, *Fkbp5*, *Fbxo31*, and *Eif4e*. X-axis categories: CTL (control), total *Mafa* mutant, muscle specific *Mafb* mutant, muscle specific *c-Maf* mutant (Sadaki et al., 2023). Statistical comparisons by one-way ANOVA with post-hoc testing. Asterisks indicate significance: *p < 0.05, **p < 0.01, ***p < 0.001.

**Supplementary Figure 11.**
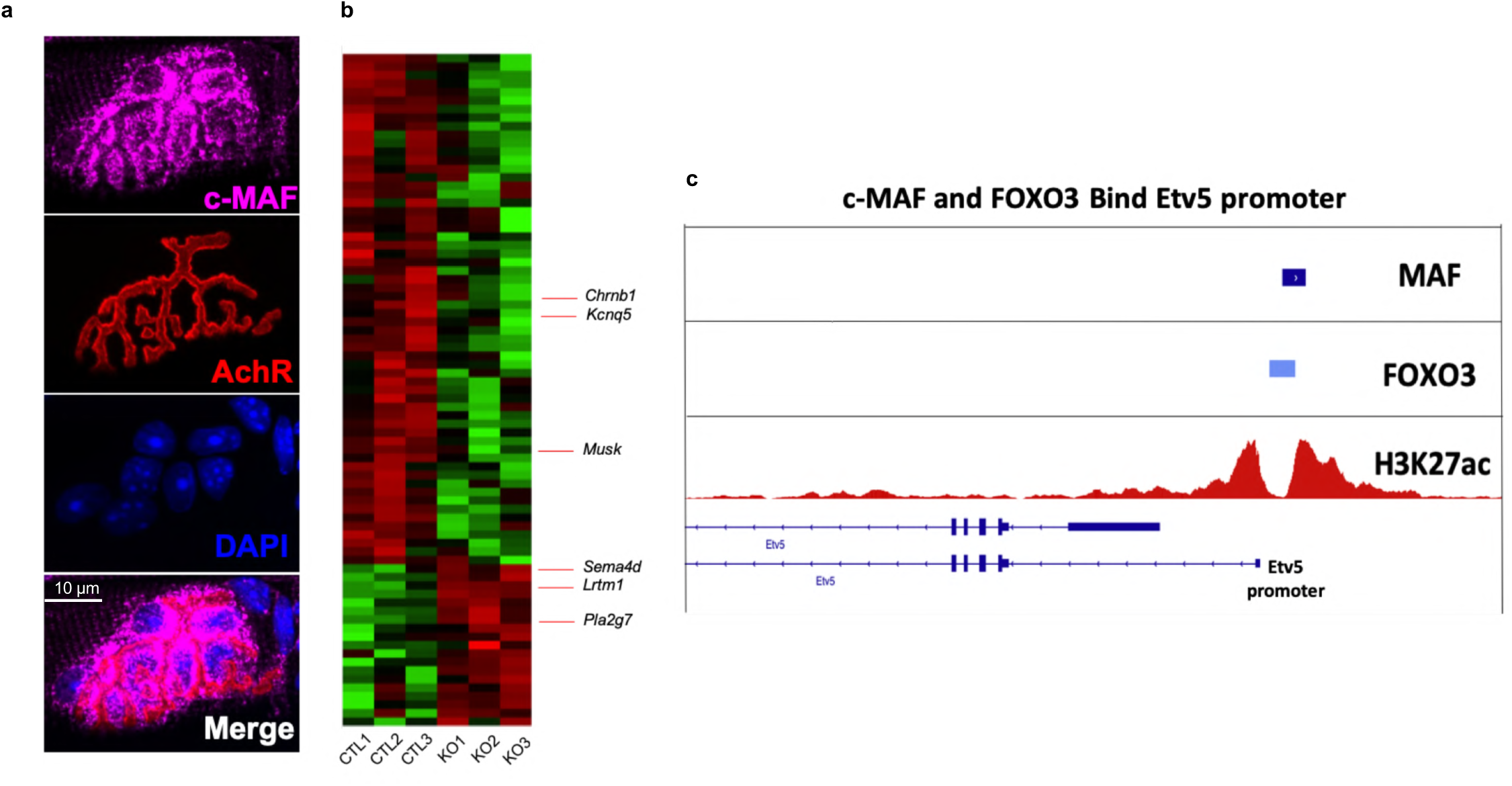
c-MAF localized to NMJ myonuclei and regulated *Etv5* expression through direct chromatin binding. **a**, Immunofluorescence of c-MAF localization at neuromuscular junctions. Four panels showing: c-MAF (magenta), α-Bungarotoxin (red), DAPI (blue), and merged image. c-MAF signal concentrated in nuclei clustered at NMJ postsynaptic region, colocalizing with DAPI-positive myonuclei adjacent to AChR-positive endplate. Confocal maximum intensity projection. Scale bar represents 10 µm. **b**, Heatmap showing expression of NMJ-enriched genes dysregulated in *c-Maf* mutant muscle. Rows representing genes identified from snRNA-seq as enriched at NMJ and differentially expressed between control (CTL) and *c-Maf* mutant conditions. Columns showing individual samples. Color scale: red indicates low expression (downregulated), green indicates high expression (upregulated), black indicates intermediate levels. Key genes labeled on right margin include *Chrnd1*, *Kcnq5*, *Musk*, *Sema4d*, *Lrtm1*, and *Pla2g7*. **c**, ChIP-seq coverage tracks at *Etv5* gene locus showing c-MAF and FOXO3 binding (Peng et al., 2017). Top two tracks show binding site positions for MAF (dark blue bar) and FOXO3 (light blue bar) within the promoter region of *Etv5*. Third track displays H3K27ac histone modification enrichment (red peaks) indicating active chromatin at *Etv5* promoter region. Bottom track showed *Etv5* gene structure with exons (thick blue boxes) connected by introns.

**Supplementary Figure 12.**
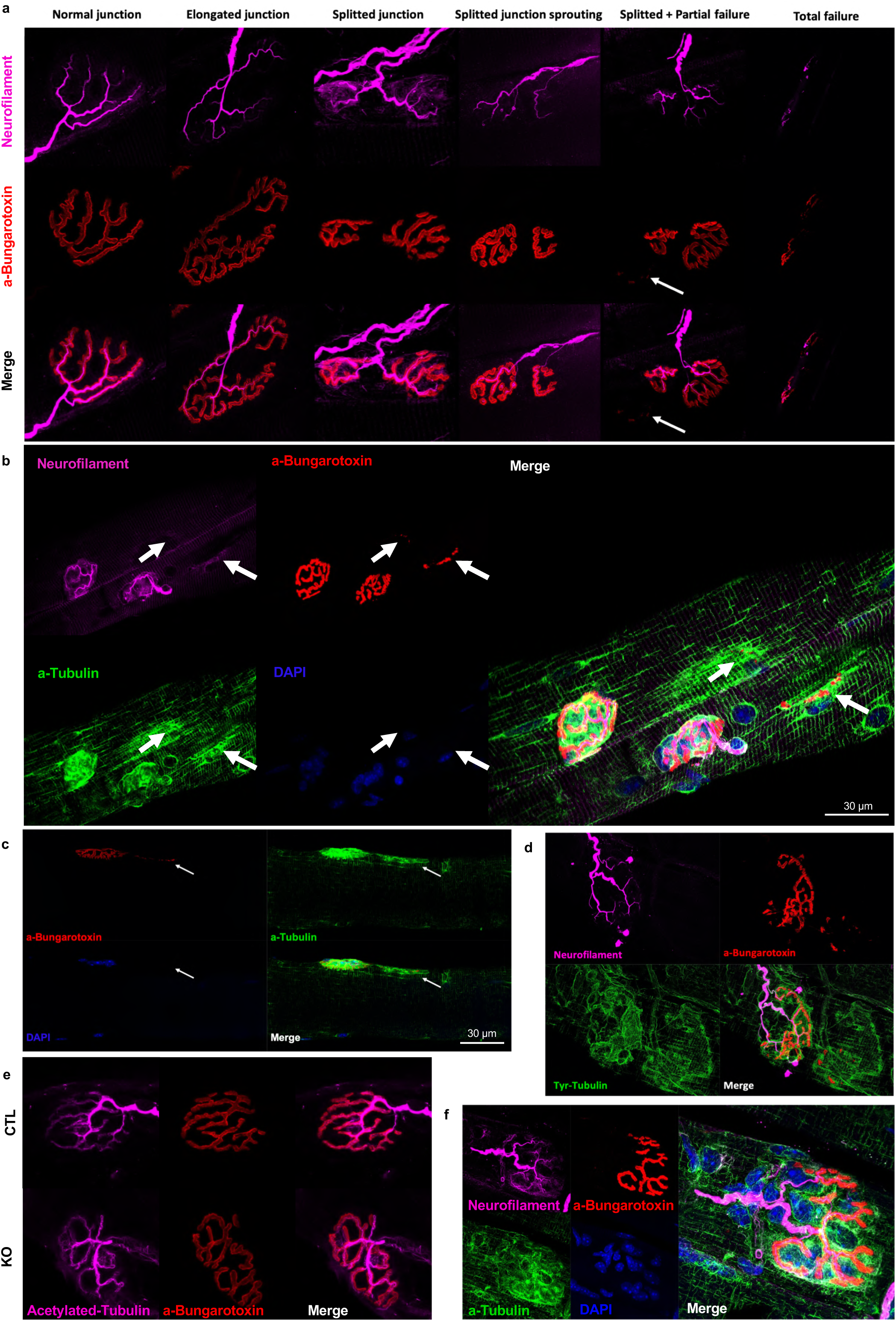
Progressive NMJ fragmentation with presynaptic microtubule disorganization in *c-Maf* mutant. **a**, Classification of NMJ morphology phenotypes in *c-Maf* mutant myofibers. Three rows show Neurofilament (magenta, presynaptic), α-Bungarotoxin (red, AChR), and merged images across six categories: normal junction, elongated junction, splitted junction, splitted with sprouting, splitted with partial denervation, and total denervation. White arrows indicate contact failure sites showing progressive fragmentation. **b**, Multi-channel immunofluorescence of NMJ architecture of *c-Maf* mutant. Panels show Neurofilament (magenta), α-Bungarotoxin (red), α-Tubulin (green), DAPI (blue), and merged image demonstrating spatial relationships between presynaptic terminal, postsynaptic membrane, and microtubule cytoskeleton. White arrows indicate ectopic AchR localization with elevated α-Tubulin staining. **c**, Ectopic AchR NMJs of *c-Maf* mutant showing α-Bungarotoxin (red), α-Tubulin (green), DAPI (blue), and merge. White arrows indicate AChR-positive postsynaptic areas with enriched microtubule organization. **d**, Ectopic AchR NMJ of *c-Maf* mutant displaying Neurofilament (magenta), α-Bungarotoxin (red), Tyr-Tubulin (green), and merge showing spatial organization of presynaptic terminal, postsynaptic membrane, and microtubule network. **e**, Sequential confocal optical sections comparing NMJs of control (CTL) and *c-Maf* mutant. Panels show Acetylated-Tubulin (magenta) and α-Bungarotoxin (red). Images demonstrating the disorganization of acetylated presynaptic Tubulin in *c-Maf* mutant (KO) compared to organized parallel arrangement in CTL, revealing pre-synaptic cytoskeletal disruption. **f**, Confocal image of multi-channel immunofluorescence showing Neurofilament (magenta), α-Bungarotoxin (red), α-Tubulin (green), DAPI (blue), and merge of *c-Maf* mutant NMJ.

**Supplementary Figure 13.**
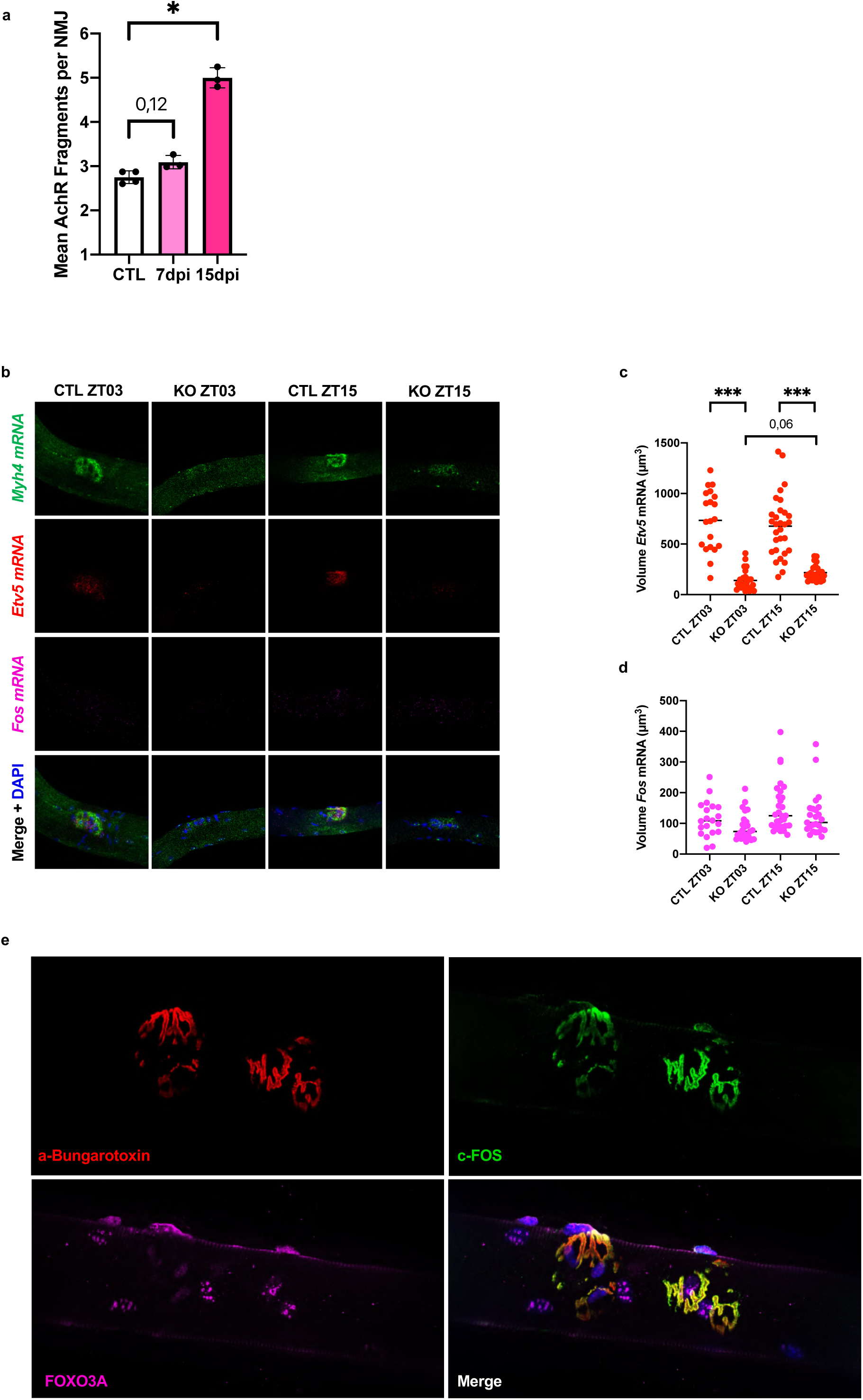
Temporal dynamics of *Etv5* and *Fos* expression at neuromuscular junctions following muscle-specific *c-Maf* deletion. **a,** Quantification of acetylcholine receptor (AChR) fragmentation at neuromuscular junctions in control (CTL), 7 days post-tamoxifen injection (7dpi), and 15 days post-tamoxifen injection (15dpi) in *HSA-CreERT2; c-Maf* mutant tibialis anterior muscle. Progressive NMJ fragmentation occurs following *c-Maf* deletion, with significant increases in mean AChR fragment number by 15 days post-tamoxifen. **b**, Representative fluorescence *in situ* hybridization images showing *Myh4* (green), *Etv5* (red), and *Fos* (magenta) mRNA expression with DAPI nuclear counterstain (blue) in TA isolated fibers of *c-Maf* inducible mutant 4 weeks after tamoxifen injection at Zeitgeber time 3 (ZT03) and Zeitgeber time 15 (ZT15). **c**, Quantification of *Etv5* mRNA puncta volume at NMJ showing significant reduction in mutant animals at both ZT03 and ZT15 compared to CTL (***P < 0.001), and a trend toward augmentation in KO ZT15 versus KO ZT03 (P = 0.06). Each dot represents a single measurement; horizontal lines indicate mean values. Statistical test: Two-way ANOVA. **d**, Quantification of *Fos* mRNA puncta volume at the NMJ across genotypes and Zeitgeber times. No significant differences were observed between groups. Each dot represents a single measurement; horizontal lines indicate mean values. **e,** Representative confocal z-stack maximum projection images of neuromuscular junctions at 4 weeks post-tamoxifen injection showing co-localization of α-bungarotoxin (α-BTX, red) labeling postsynaptic AChRs, c-FOS (green), and FOXO3A (magenta) indicating activation of atrophy-related transcriptional programs. Merge image shows nuclear DAPI (blue) with overlay of all channels, demonstrating nuclear accumulation of FOXO3A in myonuclei associated with fragmenting neuromuscular junctions following *c-Maf* deletion.

